# Microsaccadic eye movement generation is sufficient to enhance peripheral visual sensitivity

**DOI:** 10.1101/2024.12.23.630151

**Authors:** Tong Zhang, Xiaoguang Tian, Tatiana Malevich, Matthias P. Baumann, Ziad M. Hafed

**Author notes:** **Correspondence:** Ziad M. Hafed, Otfried-Müller Str. 25, Tübingen, 72076, Germany. Contributed equally (Tong Zhang, Xiaoguang Tian).

## Abstract

Microsaccades have been convincingly linked to extrafoveal covert attention shifts for more than two decades. However, the direction of causality between individual microsaccade generation and an alteration in both extrafoveal visual sensitivity and behavior remains debated: do microsaccades merely reflect, perhaps probabilistically, an altered extrafoveal sensitivity, or is the act of generating microsaccades sufficient, on its own, to modify such sensitivity? Using a novel exploitation of real-time retinal image stabilization, behavior, and neurophysiology in the superior colliculus of rhesus macaque monkeys, we show that exclusive experimental control over foveal oculomotor state is entirely sufficient to influence extrafoveal sensitivity. This happens for eccentricities as large as ∼50 times those associated with microsaccades, and it also takes place in the absence of any differential attentional demands. Most importantly, such influence is mediated through well-known, classic pre- and post-saccadic visual processing changes. Thus, seemingly-innocuous subliminal eye movements do constitute an integral component of cognitive processes like attention.

## Introduction

Microsaccade directions were discovered to be correlated with the directions of extrafoveal covert visual attention shifts almost one quarter of a century ago (1, 2). These results were replicated many times (3, 4), including in monkeys (5, 6, 7, 8, 9, 10, 11, 12, 13, 14), and they were also extended to the allocation of attention towards items recently committed into working memory (15, 16, 17, 18).

At first glance, a correlation between microsaccades and covert visual attention seems entirely plausible, especially given the long history (19, 20, 21, 22) of studies linking attentional processing to oculomotor programming. However, further contemplation over this correlation reveals a deeper and fundamental question in cognitive neuroscience, related to the direction of causality. For example, in 2002, Hafed and Clark (1) checked extrafoveal perceptual performance when their discrimination stimuli appeared in the absence of nearby microsaccades; their differential task performance effects were lost, and they wrote: “suggesting that attention may not have shifted during those trials” (1). This observation went largely dormant for another ten years, until Hafed and colleagues discovered a neuronal mechanism for microsaccade generation in the superior colliculus (SC) (23). This discovery, underscoring the high mechanistic similarity between microsaccades and larger saccades (24), now raised the possibility that individual microsaccade generation can alter, perhaps through corollary discharge (25, 26, 27, 28), visual processing during peri-movement epochs, exactly like larger saccades do (29). Armed with this, Hafed revisited the 2002 mystery in 2013, and indeed found that covert attentional enhancement effects, in cueing paradigms invoking shifts of attention, were essentially only contingent on the appearance of discrimination stimuli during pre-microsaccadic epochs, and with specific directions relative to microsaccade directions (30). These results were further supported by several extensive experiments (with or without explicit attentional tasks) involving human and monkey behavior, monkey neurophysiology, and computational models (4, 8, 31, 32, 33).

More recently, although without explaining it using as clear a mechanistic process as the one of peri-saccadic visual alterations, it was reported that cortical neuronal modulations in a classic covert attentional task depended entirely and exclusively on microsaccades (13). This publication has garnered much more debate (11, 12, 16, 17, 18) than in the case of Hafed’s peri-saccadic hypothesis, with the net result being that the overall causality question of microsaccades is still unresolved.

Here, we aimed to causally test a role for microsaccade generation in altering extrafoveal visual sensitivity, even when attentional state is explicitly controlled for. We were motivated by Hafed’s original pre-microsaccadic enhancement phenomenon. In this phenomenon, if an extrafoveal visual stimulus, like discrimination stimuli in classic covert attentional tasks (4), appears right before a given microsaccade, then visual sensitivity and subsequent behavioral responses are enhanced (4, 31, 32, 33, 34). Moreover, such enhancement is qualitatively similar to known attention-related enhancements during cueing paradigms driving attention shifts (35, 36, 37, 38, 39). Even though our previous work already showed that enhanced visual sensitivity could happen for any microsaccade, including in the absence of attentional tasks (31, 34), this work still did not establish a potential direction of causality: it could still be that extrafoveal visual sensitivity is independently enhanced (due to any factor other than microsaccade generation), and that it is this enhancement that triggers a short-latency microsaccade (hence, the appearance of pre-microsaccadic enhancement). To explicitly test this, here we used a paradigm allowing exclusive experimental control over only foveal, but not extrafoveal, oculomotor and neuronal states. This paradigm exploits the fact that microsaccades fundamentally serve the function of optimizing eye position on the foveal target (32, 33, 40, 41, 42), even in the presence of small head movements during real world scenarios (43). In what follows, we show that this was sufficient to significantly modulate extrafoveal visual sensitivity, and we also show human perceptual implications of this. Then, we discuss how our evidence, coupled with accumulated knowledge about the underlying neurobiology of the oculomotor system during exogenous and endogenous stimulation (10, 44, 45, 46, 47), can allow easily resolving many of the above debates about microsaccades and attention.

## Results

We established a causal role for foveal oculomotor state, including microsaccade generation, in influencing extrafoveal visual sensitivity and behavior, and independently of the allocation of covert visual attention. To do so, we designed (32, 33, 48) a novel real-time, gaze-contingent retinal image stabilization paradigm explicitly forcing a tiny, but constant, foveal visual error during fixation. Prior research (32, 33, 40, 41) has shown that such a constant foveal visual error should robustly modulate microsaccade directions, attesting to the role of microsaccades in optimizing eye position (40, 41, 43). Here, we exploited this technique by pairing it with a task requiring visually-driven behavioral responses to extrafoveal stimuli. In what follows, we show that our experimental manipulation: 1) allowed exquisite control over microsaccade directions while maintaining task-related attentional state constant; 2) that it allowed exclusive control over only the foveal, but not extrafoveal, SC neuronal state before extrafoveal visual stimulus onset; and 3) that exclusively manipulating foveal oculomotor state (both behaviorally and neuronally) was adequate to modulate extrafoveal visually-driven neuronal sensitivity and behavior. We then conclude with human perceptual experiments demonstrating direct homologs of our neurophysiological results, revealing that foveal action is indeed sufficient to modulate extrafoveal vision.

### Experimental control over foveal oculomotor state

Our monkey paradigm involved maintaining prolonged gaze fixation while monitoring a single extrafoveal location at which a visual stimulus was to appear. Once the extrafoveal stimulus appeared, the monkeys reflexively foveated it as quickly as possible (Methods). Throughout all trials within a session, the extrafoveal visual stimulus location was constant, allowing us to control the animals’ extrafoveal attentional states across the different foveal conditions that we tested. These foveal conditions, randomly interleaved, included either a stable fixation spot on the display (control), or a spot that was synchronously moved with instantaneous eye position to minimize its motion on the retina (Methods). Critically, in the trials with foveal retinal image stabilization, we forced a constant offset between instantaneous eye position and the displayed fixation spot position such that there was always a tiny (∼3.5-7.7 min arc; Methods; S1 Figure) visual error for as long as retinal image stabilization was applied (Fig. 1A). Such a constant foveal visual error generated a “perpetual” microsaccade motor plan; it was thus expected (49, 50, 51, 52, 53, 54) to elevate low-level foveal SC activity congruent with the visual error metrics. Moreover, we chose the direction of the forced foveal visual error to either be the same (Fig. 1A, blue), opposite (Fig. 1A, red), or orthogonal (Fig. 1A, cyan and green) to the direction of the upcoming extrafoveal visual stimulus onset at the end of the fixation epoch (Fig. 1A, S1 Figure) (Methods).

**Figure 1.**
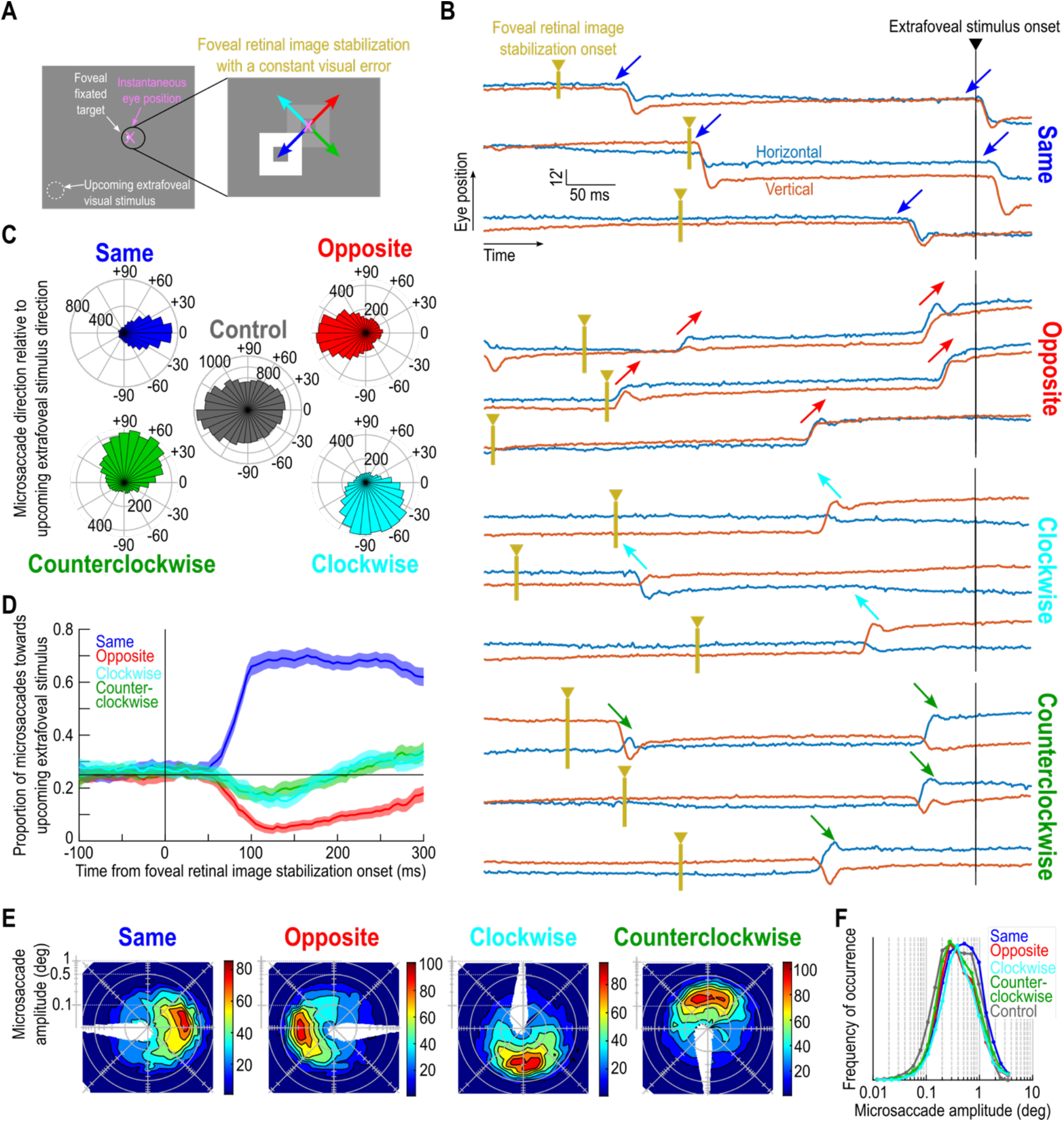
Foveal retinal image stabilization allows exquisite control over microsaccade directions. **(A)** Four monkeys fixated a small spot in anticipation of an extrafoveal visual stimulus onset. During fixation, we forced a tiny, but constant, visual error between the displayed spot location (white in the magnified inset) and the instantaneous eye position (magenta) (Methods). Across trials, the visual error could be in different directions relative to the direction of the upcoming extrafoveal stimulus. **(B)** Horizontal and vertical eye position traces from 12 example trials (monkey N). We grouped the trials according to the direction of foveal visual error, and we aligned the trials to the onset of the extrafoveal visual stimulus (placed in the lower left quadrant in this session). Small colored arrows indicate the direction of the corresponding microsaccade. During retinal image stabilization, each triggered microsaccade was in the direction of the forced visual error. **(C)** Distributions of microsaccade directions (relative to the direction of the upcoming extrafoveal visual stimulus) for all movements occurring during retinal image stabilization (colored histograms). Data across all sessions and monkeys are shown. The gray histogram shows the distribution in control trials with no foveal retinal image stabilization. **(D)** Microsaccade directions were dominated by foveal retinal image stabilization as soon as it was applied. Within approximately 50-60 ms from foveal retinal image stabilization onset, microsaccade directions relative to the extrafoveal visual stimulus direction reflected the applied foveal visual error. Error bars denote 95% confidence intervals, and 0.25 on the y-axis indicates unbiased microsaccade directions. **(E)** Microsaccade amplitudes (on a log-polar scale (24, 52, 57)) as a function of microsaccade directions in the different foveal retinal image stabilization conditions (z-axis indicates microsaccade counts per shown bin). Foveal retinal image stabilization dictated microsaccade directions, while keeping microsaccade amplitude distributions largely consistent with the forced visual error amplitudes that we imposed. **(F)** Same as **E** but pooling across microsaccade directions. S2 Figure show individual monkey results, and S3 Figure shows additional properties of microsaccades that emerged as a result of retinal image stabilization.

In all four tested monkeys, our paradigm succeeded in experimentally dictating microsaccade directions leading up to, and around, the onset of the extrafoveal visual stimulus onset. Consider, for example, the eye position traces in Fig. 1B. Each horizontal/vertical pair represents a single sample trial, and the trials are grouped according to the direction of forced foveal visual error. We aligned all traces to the onset of the extrafoveal visual stimulus, which was the end of the fixation epoch (Methods). The onset of foveal retinal image stabilization on each trial is also indicated. Since the extrafoveal visual stimulus location in this example session was in the bottom left quadrant, a forced foveal visual error in the same direction as the extrafoveal stimulus caused downward leftward microsaccades (top three example trials, with a small blue arrow placed above each detected microsaccade indicating its direction for easy visualization). All microsaccades were towards the bottom left quadrant. Note also that the first two trials additionally had the stimulus onset occur right before a microsaccade; we will revisit this special case (30, 31, 34) of an immediate pre-microsaccadic extrafoveal visual stimulus event later below. The remaining three groups of example trials in Fig. 1B (from the same session) show that when forced foveal visual error was opposite the direction of the upcoming extrafoveal visual stimulus, all microsaccades were upward/rightward (red); for orthogonal forced foveal visual errors, the microsaccade directions reflected the direction of the experimental manipulation (cyan and green). Thus, foveal retinal image stabilization strongly dictated microsaccade directions during fixation, consistent with our earlier behavioral demonstrations of this in monkeys (32, 33).

Across all sessions and monkeys, we collected all microsaccades happening during foveal retinal image stabilization, and we plotted their direction distributions relative to the direction of the upcoming extrafoveal visual stimulus (colored histograms in Fig. 1C). Microsaccade directions were overwhelmingly dictated by the direction of forced foveal visual error: when it was in the same direction as the extrafoveal visual stimulus, most microsaccades had an angular difference of near zero from the direction of the extrafoveal stimulus (blue); when it was opposite, the microsaccades were mostly 180 deg away from the extrafoveal visual stimulus direction (red); and when it was orthogonal, the movements were mostly orthogonal (cyan/green). In contrast, microsaccade directions were random with respect to the extrafoveal stimulus direction in the control condition (gray histogram in Fig. 1C). S2 Figure shows individual monkey results.

Not only were microsaccade directions successfully experimentally controlled, but such control emerged almost instantaneously after starting retinal image stabilization (Fig. 1D). We binned microsaccade directions into four bins relative to the direction of the upcoming extrafoveal visual stimulus (Methods). Within around 50-60 ms from retinal image stabilization onset, microsaccade directions were strongly biased by the direction of the forced foveal visual error (error bars denote 95% confidence intervals). With longer delays (e.g. the red curve in Fig. 1D), there was a small trend for this effect to weaken. This was expected because the forced visual error displaced eye position from the display center; the longer that this happened, the more likely were the monkeys to try to fight this, in order to not lose the possibility of reward if the eye moved so far out as to prevent the extrafoveal visual stimulus from being within the display boundaries when it appeared. Such “fighting” is not too different from the complementary, and thus sometimes competitive, interaction that exists between drifts and microsaccades during normal fixation without retinal image stabilization (55, 56); it was merely amplified when we controlled the direction of the foveal visual error for a prolonged period of time.

Importantly, our foveal manipulation was so subtle that it did not drastically alter microsaccade amplitude distributions (Fig. 1E, F). With large forced visual errors of more than even just 1 deg in amplitude, retinal image stabilization can massively destabilize the oculomotor control system, since the error that this system attempts to attenuate with eye movements never decreases (S1 Movie) (58); in other words, “opening” the negative feedback control loop reveals the high gain of the saccadic system (58). However, we used an extremely small visual error (Fig. 1A), and with limited retinal image stabilization durations (Methods). This allowed us to successfully maintain relatively stable gaze fixation even during retinal image stabilization (Fig. 1B), and this, in turn, resulted in small microsaccade amplitudes (Fig. 1E, F). Importantly, these amplitudes, slightly larger than amplitudes during control trials (as expected from a forced non-zero visual error on retinal image stabilization trials), were consistent (57) with what to expect from microsaccades (mean+/-SD: 28.8+/-23.4 min arc for control, 38.4+/-33 min arc for same, 31.8+/-28.8 min arc for opposite, 34.2+/-31.2 min arc for clockwise, and 33.6+/-31.2 min arc for counterclockwise). Thus, highly precise foveal retinal image stabilization (with tiny forced visual errors) allowed us exquisite experimental control over microsaccade directions, without otherwise drastically altering other fixation properties (like drifts and microsaccade amplitudes).

We also inspected additional properties of microsaccades during foveal retinal image stabilization, in order to confirm that the movements were indeed genuine outcomes of our experimental manipulation. We analyzed microsaccade rate around the time of retinal image stabilization onset (S3A Figure). Consistent with the fact that the onset resulted in a foveal sensory transient, there was a subtle short-latency reduction in microsaccade rate, followed later by a rebound. This is a classic “orienting response” of the oculomotor system, which enables it to coordinate between realizing unfinished motor plans at the time of the sensory transients and generating new responses to these transients (44). Importantly, the time of microsaccade direction biasing (Fig. 1D) coincided with the time of microsaccadic inhibition (see the inset in S3A Figure), revealing both classic components of the expected orienting response (rate modulation and direction biasing) (8). Interestingly, when we then split the microsaccade rate curves as a function of the direction of foveal retinal image stabilization, we observed an additional effect of target expectation in the post-inhibition microsaccades that we had not initially expected: the monkeys generated fewer rebound microsaccades if the foveal error was in the opposite direction from the upcoming target location than when it was in the same direction (S3B Figure). Such rebound microsaccades reflect top-down control over microsaccades (9, 42), suggesting that there was evidence of target expectation even in the monkeys’ microsaccades. Such expectation, due to the constant extrafoveal stimulus location per session, also emerged in the form of more directionally-accurate microsaccades in the same than opposite condition. For example, inspecting the direction distributions of Fig. 1C and those of the individual monkeys (S2A-D Figure) revealed systematically tighter distributions in the same condition. We confirmed this observation in S3C, D Figure by measuring the standard deviation of microsaccade directions. As we discuss further below, even the later “overt” saccadic responses to the extrafoveal stimuli revealed a component of target expectation. This prompted our next analysis of the retinal image stabilization experimental manipulation, this time from a neuronal perspective, to again underscore its utility for our current purposes despite these subtle target expectation effects.

### Exclusive experimental control over foveal SC neuronal state

We were ultimately interested in the effects of foveal oculomotor state on extrafoveal visual sensitivity and behavior. However, we first further established that our experimental manipulation was indeed exclusively limited to foveal state. We recorded from the SC’s foveal representation (52) during retinal image stabilization (Fig. 2A), and we analyzed neuronal activity when the forced foveal error was congruent (same) or incongruent (opposite) with the recorded foveal response field (RF) directions. In the latter case, the experimental manipulation generated “perpetual” foveal oculomotor commands in the other SC rather than the one that we were recording from.

**Figure 2.**
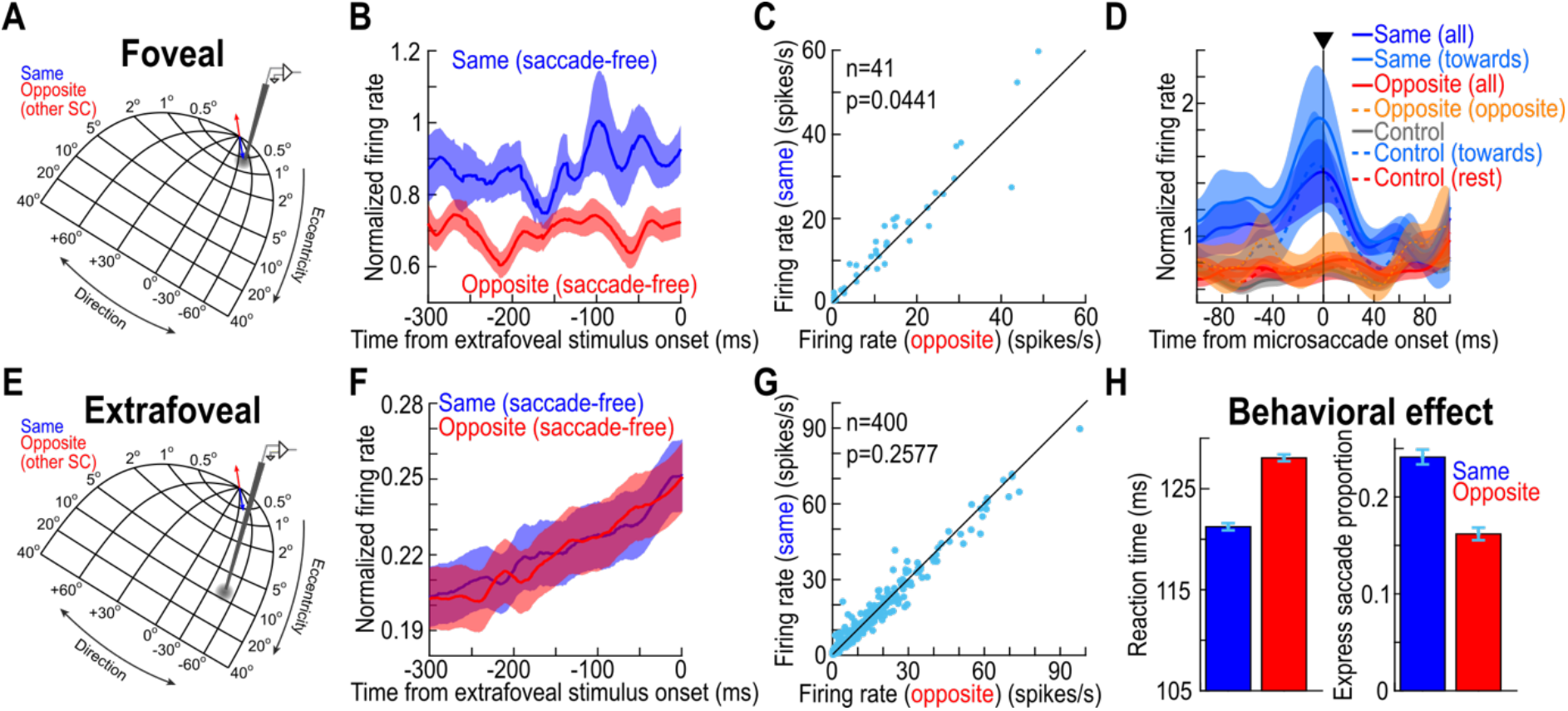
Exclusive differential modulation of foveal, but not extrafoveal, superior colliculus (SC) activity alters subsequent extrafoveal visually-driven behavior. **(A)** To confirm that foveal retinal image stabilization exclusively modulated the foveal SC, we recorded from this SC region (the schematic shows the SC’s anatomical representation of the contralateral visual field (52, 64)). A forced visual error in the same direction (blue) as the upcoming extrafoveal visual stimulus was expected to elevate foveal SC activity representing the goal location caused by the error (49, 50, 59). **(B)** Microsaccade-free (Methods) firing rate of foveal SC neurons in the final 300 ms of retinal image stabilization. When the forced visual error was in the same direction as the recorded neurons (blue), activity was elevated relative to opposite visual errors (red). S4B, C Figure shows results with orthogonal retinal image stabilization, as well as control. **(C)** Raw firing rates of all foveal neurons. Forced visual error caused a “perpetual” movement plan that elevated foveal SC activity in the appropriate direction (49, 50, 59). **(D)** During retinal image stabilization, microsaccades in the same condition (blue) were associated with expected motor bursts (23, 24) since their directions were consistent with the response fields of the recorded foveal neurons (light blue shows microsaccades in the same condition that were genuinely towards the RF direction). This did not happen for the opposite (red) and control (gray) conditions, except when congruent microsaccades did occur (e.g. dashed light blue). The orange curve further shows that microsaccades in the opposite condition that were genuinely opposite in direction from the foveal RF’s also did not elevate SC activity; thus, the slightly larger variance in microsaccade directions seen in the opposite condition (S3C, D Figure) did not corrupt our manipulation’s aim of differentially controlling foveal SC state. **(E)** Exclusive differential modulation of foveal SC state (**A**) should not affect pre-stimulus extrafoveal SC activity during fixation. **(F)** Same as **B** but for extrafoveal neurons. There was no dependence on foveal visual error. S4F Figure shows results from the orthogonal conditions as well. **(G)** Same as **C** but for extrafoveal neurons, again showing no dependence on foveal visual error. Thus, our experimental manipulation exclusively controlled fovea, but not extrafoveal, SC neuronal state. **(H)** Visually-driven extrafoveal behavior, after extrafoveal stimulus onset (Methods), clearly depended on foveal SC state, even though pre-stimulus extrafoveal SC state was unmodulated. Error bars denote SEM in the left plot and 95% confidence intervals in the right plot. The underlying data are included in S1 Data. S6 Figure show results from the orthogonal conditions.

Microsaccade-free (Methods) foveal SC activity directly reflected the forced foveal visual error direction. Specifically, whenever the forced foveal visual error was in the same direction as the recorded neurons, microsaccade-free tonic activity was elevated relative to the opposite foveal error (Fig. 2B) and control (S4A-C Figure). This is expected: foveal retinal image stabilization forced an oculomotor “goal” location slightly deviated from the current line of sight, thus resulting in a “perpetual” microsaccade plan; such a plan slightly elevated low-level firing activity in the foveal SC in the absence of microsaccades (23, 49, 50, 53, 59) (Fig. 2C). Indeed, the elevation that we observed in the same condition was quantitatively not too different from previous observations of foveal SC goal representations (e.g. (50)), which is remarkable given how small the foveal errors that we introduced (only 3.5-7.7 min arc) were relative to foveal SC RF sizes (52). We also included the orthogonal conditions of Fig. 1, which (due to population coding (52, 60)) always expectedly gave intermediate results between those observed for the same and opposite conditions (S4C Figure).

More importantly, when microsaccades did occur, the effect of forced visual error direction (on the foveal SC’s activity) was strongly amplified, exactly as planned (Fig. 2D): around any given microsaccade, there was a strong motor burst if the microsaccade was directed towards the recorded foveal RF’s. This happened in the same condition (Fig. 2D, blue) since the majority of microsaccades were, by virtue of experimental control (Fig. 1B-E), congruent with the foveal RF locations (Fig. 1), but it did not happen for the opposite (Fig. 2D, red) and control (Fig. 2D, gray) conditions. To convince ourselves that our experimental manipulation was indeed primarily tapping into dictating microsaccade directions without drastically changing other properties (Fig. 1), we then separated the microsaccades in the control condition according to whether they were directionally congruent with the recorded foveal RF locations or not (Fig. 2D, dashed) (Methods). Whenever microsaccades during control had the same direction as the foveal RF’s, we obtained expected (23, 24, 52) microsaccade-related motor bursts as well. Figure 2D also shows the microsaccades in the same condition that were genuinely congruent with the neurons’ RF’s; since they were the majority of the microsaccades anyway (Fig. 1), the neurons also strongly burst for these movements. Similarly, microsaccades in the opposite condition that were genuinely opposite the recorded RF’s did not elevate SC activity (orange), allowing us to control for the slightly larger variance of directions in opposite microsaccades seen in S3C, D Figure. Thus, foveal retinal image stabilization not only dictated microsaccade directions, but it also differentially modulated the state of the foveal SC, in a manner consistent with known oculomotor programming processes.

Most critically, differential modulation of foveal SC state did not “leak” into the extrafoveal SC during fixation. We repeated the same pre-stimulus analyses above, but this time for all of our 400 extrafoveal SC neurons (Fig. 2E), spanning all four quadrants of visual space (S5 Figure). During retinal image stabilization, the direction of the forced foveal visual error did not differentially alter extrafoveal SC pre-stimulus activity at all (Fig. 2F, G). This was also true for the orthogonal visual error conditions (S4F Figure). Note that pre-stimulus activity did tend to slowly rise (equally for the different retinal image stabilization conditions; Fig. 2F, S4E, F Figure) in anticipation of extrafoveal stimulus onset; this is a known property of SC neurons with increasing odds of target onset (61, 62, 63), and it was visually amplified by our normalization procedure (Methods; see, for example, Fig. 3 with the raw measurements and much less visible pre-stimulus elevation). Indeed, this is yet another form of evidence that some aspect of target expectation and arousal was triggered by our foveal retinal image stabilization experimental manipulation (similar to the secondary microsaccadic effects seen in S3B-D Figure). However, it was independent of the differential modulation of the foveal SC state, which is what we aimed to achieve.

**Figure 3.**
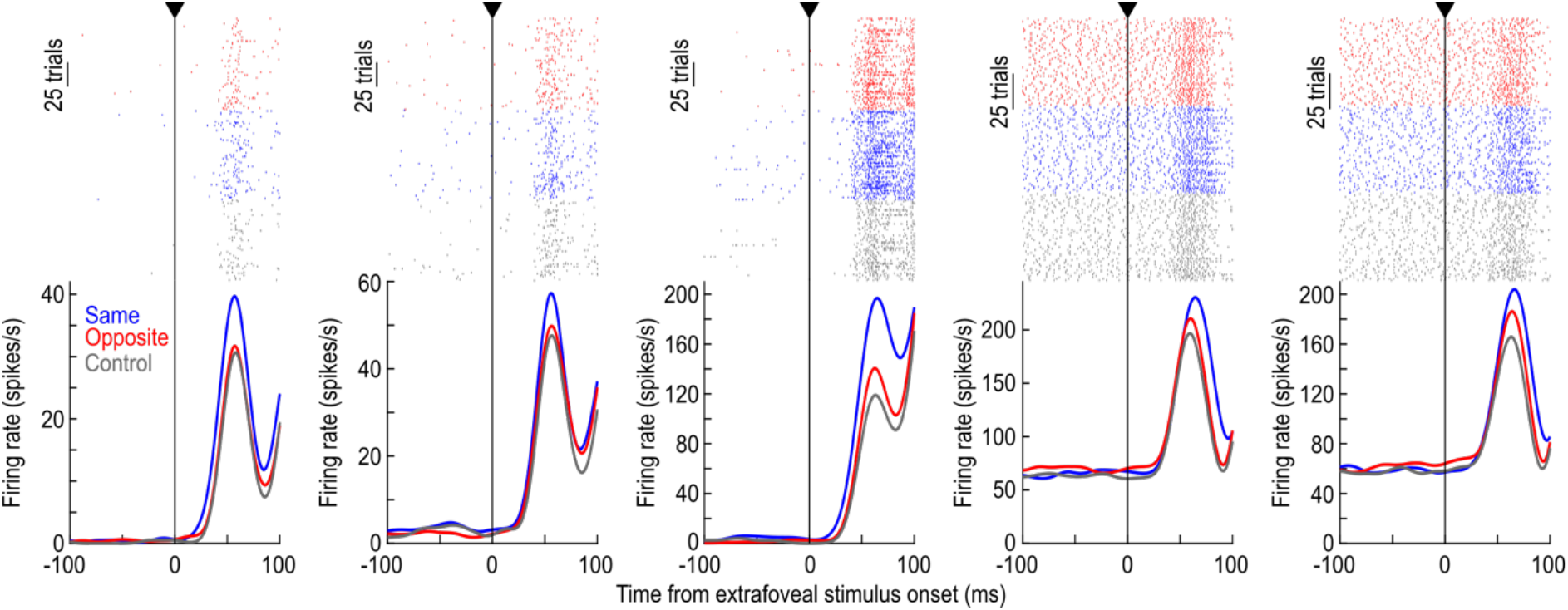
Extrafoveal visual sensitivity is differentially modulated by exclusive control over pre-stimulus foveal SC state. Each panel shows the visual responses of an individual example neuron (the first three are from monkey N, and the last two are from monkey A) under three different foveal visual error conditions: control (without foveal retinal image stabilization), same (forced foveal visual error in the direction of the upcoming extrafoveal visual stimulus), and opposite (forced foveal visual error opposite the direction of the upcoming extrafoveal visual stimulus). Bottom shows firing rate curves, and top shows individual trial spike time rasters (grouped according to condition). All neurons exhibited the strongest visual bursts for the same condition. The opposite condition also showed enhanced visual bursts relative to control, likely due to motor preparation (S4 Figure; also see S3B-D Figure), but the differential between the same and opposite conditions was always in favor of the same condition (consistent with the behavioral effects of Fig. 2H). Note how pre-stimulus activity in all neurons was not affected by the fact that there was forced foveal retinal image stabilization in the same and opposite conditions. This is consistent with Fig. 2E-G, S4D-F Figure, and suggests that the differentially modulated foveal SC state specifically differentially modulated extrafoveal visually-evoked sensitivity and not baseline firing rate patterns. Numbers of trials per condition are evident from the rasters. Orthogonal retinal image stabilization conditions always gave intermediate results between the same and opposite conditions (also see S7-S9 Figures).

Thus, before extrafoveal visual stimulus onset, our directional experimental manipulation exclusively differentially modulated foveal (Fig. 2A-D), but not extrafoveal (Fig. 2E-G), SC neuronal state. We next show how this exclusivity was still sufficient to not only differentially modulate extrafoveal behavior (Fig. 2H), but to also differentially alter visually-evoked extrafoveal neuronal sensitivity.

### Differential modulation of extrafoveal visual sensitivity

Behaviorally, as soon as the extrafoveal stimulus appeared, the monkeys foveated it faster when pre-stimulus foveal SC state (Fig. 2A-D) was elevated than when it was not (Fig. 2H, left). The frequency of express saccades (65) (Methods) was also higher for the same condition (Fig. 2H, right). Given past work on the links between SC visual neuronal sensitivity and reaction times (66, 67, 68, 69, 70, 71, 72, 73), we hypothesized that visually-evoked SC bursts in our paradigm must have been modulated by pre-stimulus foveal retinal image stabilization; otherwise, the behavioral results of Fig. 2H would not have come about. This was indeed the case, as can be seen from the five example extrafoveal SC neurons of Fig. 3 (the first three neurons are from monkey N; and the next two neurons are from monkey A). In each case, visually-evoked activity was the highest for the same condition, even though pre-stimulus activity was similar across all foveal state conditions (consistent with Fig. 2F, G). Thus, foveal action was sufficient to modulate extrafoveal visual sensitivity, as we previously predicted (1, 4, 30, 31, 42). And, it did so without differentially altering pre-stimulus extrafoveal activity.

Note also that the opposite condition in Fig. 3 still experienced some enhanced sensitivity relative to control. This was because retinal image stabilization onset was already a temporal cue to the upcoming extrafoveal stimulus onset (also see S3A Figure), and also because (by task design) the extrafoveal stimulus location was unambiguous (Methods; also see S3B-D Figure). Indeed, pre-stimulus extrafoveal SC activity was elevated ever so slightly more during all retinal image stabilization conditions than during control (S4E, F Figure) (but with no difference among the different retinal image stabilization manipulations), and reaction times were generally slowest in control trials (S6 Figure). Nonetheless, the differential effects between same and opposite conditions in Fig. 3 were only paralleled by a differential modulation in foveal, but not extrafoveal, pre-stimulus SC state (Fig. 2A-G). Thus, our results so far suggest that foveal retinal image stabilization exclusively differentially modulated only foveal pre-stimulus neuronal state (Fig. 2A-G), and that this expectedly dictated microsaccade directions (Fig. 1); most importantly, this was sufficient to also differentially modulate extrafoveal visual sensitivity (Fig. 3), and, in turn, behavior (Fig. 2H).

The above observations were consistent across extrafoveal neurons. Figure 4A shows the average population visual responses of all visual-motor neurons in our database. In this figure, we first subtracted pre-stimulus baseline activity to focus on visually-evoked bursts (Methods). Extrafoveal visual sensitivity was systematically enhanced by foveal retinal image stabilization, and it was differentially modulated across forced foveal visual error directions (compare blue and red curves). This can also be seen from the raw firing rates of Fig. 4B, C, with each panel comparing visual burst strength in the retinal image stabilization trials (same for Fig. 4B and opposite for Fig. 4C) to visual burst strength in control trials. Foveal retinal image stabilization enhanced visual sensitivity, and it did so more strongly in the same condition (greater deviations from the unity slope line for the same than opposite conditions). Quantitatively, we computed a neuronal modulation index relative to control (Fig. 4D; Methods). This index was 3.4% (mean) for the same condition and 1.5% for the opposite condition (p=4.6826×10^-5^; paired t-test comparing same and opposite modulation indices), suggesting a differential modulation of extrafoveal visual sensitivity by exclusive experimental control over foveal oculomotor state. Interestingly, visual-only neurons were not differentially modulated (S7 Figure), suggesting that our effects were specific to visual-motor SC layers.

**Figure 4.**
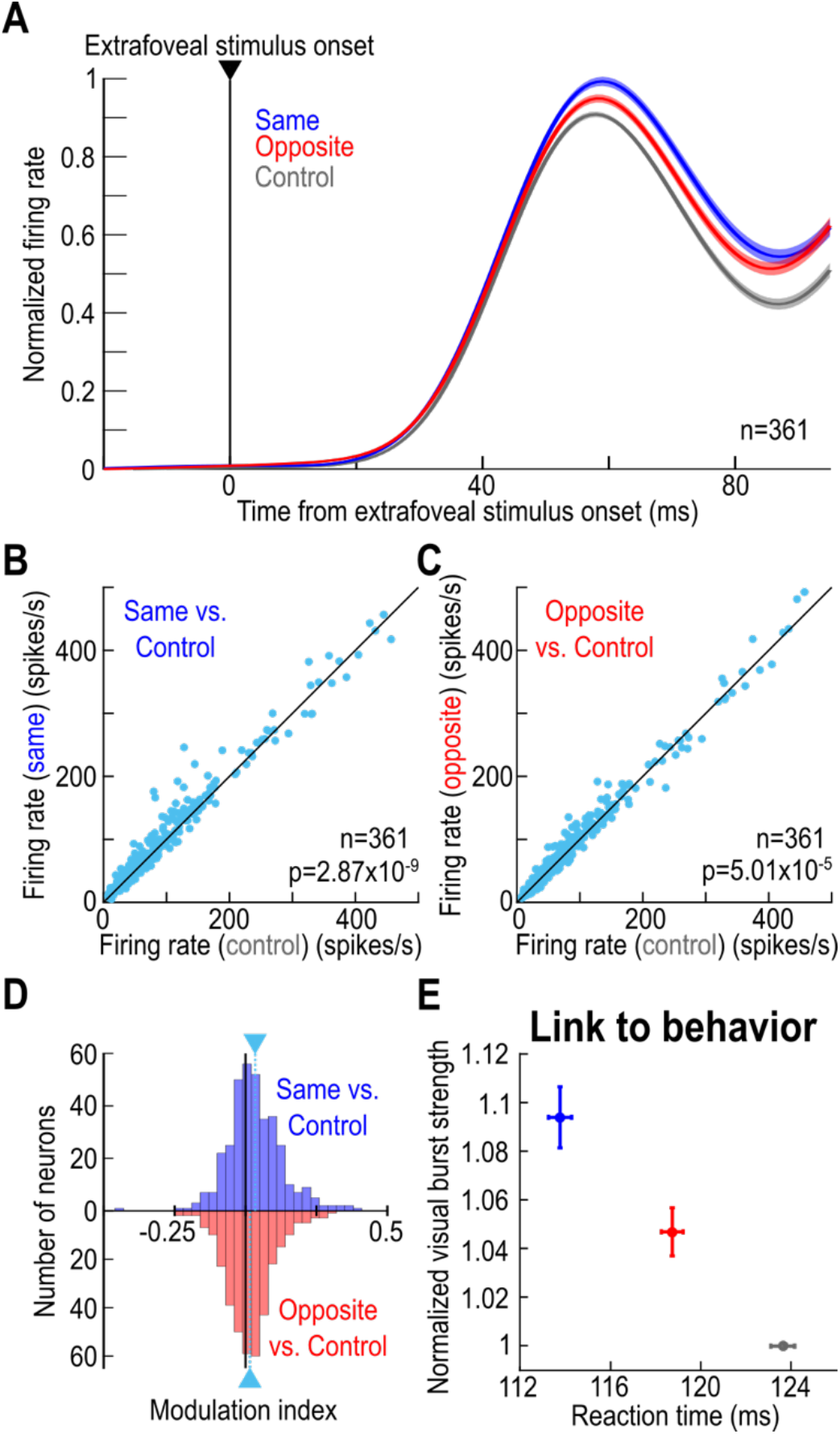
Extrafoveal visual sensitivity is differentially modulated by exclusive control over the pre-stimulus foveal SC state, and this is directly reflected in visually-driven behavior. **(A)** Population summary of the results of Fig. 3. This panel shows the average (across all visual-motor neurons) of normalized, baseline-subtracted firing rates (Methods) in the control, same, and opposite conditions (error bars denote SEM). Consistent with the example neurons, forced foveal visual error in the same direction as the upcoming extrafoveal visual stimulus was associated with the highest visual sensitivity enhancement. This happened without any “attentional” modulation, since the extrafoveal visual stimulus always appeared at the very same location across all trials. **(B)** Raw visual burst firing rates (Methods) in the same (y-axis) and control (x-axis) conditions of all visual-motor neurons. Extrafoveal visual sensitivity was enhanced with forced foveal visual error in the direction of the upcoming extrafoveal visual stimulus (paired t-test). **(C)** Same as **B** but with the opposite condition illustrated on the y-axis. There was still sensitivity enhancement, but of smaller magnitude than in **B** (also see **D** for a direct comparison). **(D)** Neuronal modulation indices (relative to control) for the same and opposite conditions (Methods). Dashed vertical lines indicate the mean across each histogram. Foveal retinal image stabilization had a stronger impact on sensitivity enhancement in the same than opposite condition. Also see Fig. 5 for when these effects were particularly amplified in the trials. **(E)** Visual sensitivity enhancement was directly (inversely) correlated with behavioral reaction times across sessions. Thus, forced foveal visual error (Fig. 1) caused exclusive differential modulation of pre-stimulus foveal (Fig. 2A-D), but not extrafoveal (Fig. 2E-G), SC neuronal state, which was sufficient to predictably (Fig. 2H and this panel) alter extrafoveal visually-driven behavior. Error bars denote SEM. The underlying data are included in S1 Data. S8 Figure shows individual monkey results, and S9 Figure shows how orthogonal foveal visual errors resulted in intermediate behavioral and neuronal effects.

The population results of Fig. 4A-D were also directly linked to the behavioral effects of Fig. 2H. Specifically, whenever visual burst strength was larger, behavioral reaction times were faster (Fig. 4E), as expected from the previous literature (66, 67, 68, 69, 70, 71, 72, 73). The difference here is that this effect was mediated by exclusive experimental control over foveal SC neuronal state before stimulus onset. Thus, foveal oculomotor state was sufficient to affect reaction times via its known impacts on extrafoveal visual sensitivity. Moreover, these results were robust at the individual monkey level (S8 Figure), which is consistent with the fact that some of these same monkeys also exhibited pre-microsaccadic sensitivity enhancements in other experimental contexts (monkey N: Chen et al., 2015 (31); monkeys A and M: Wu and Hafed, 2025 (34)). Orthogonal foveal retinal image stabilization also gave intermediate results between the same and opposite conditions (S9 Figure), as expected from SC population coding (52, 60). These observations are all consistent with our earlier findings (30, 31) that the impacts of microsaccades on extrafoveal visual sensitivity and behavior are directionally specific.

### Mediation through peri-microsaccadic impacts on extrafoveal vision

Our results demonstrate that foveal oculomotor state is sufficient to differentially modulate extrafoveal visual sensitivity. To better understand how this happens, we were motivated by our earlier idea (29) that microsaccade generation, just like larger saccade generation (74, 75, 76, 77, 78, 79), might be associated with distinct peri-movement changes in visual processing. These movement-related changes, at the level of the SC, involve both pre-movement enhanced visual sensitivity (31, 34) and post-movement reduced visual sensitivity (31, 34, 80). However, especially for pre-movement enhancement, it was always a question of whether microsaccades reflected enhanced extrafoveal sensitivity (caused by other sources) or whether the act of microsaccade generation itself was sufficient. In our case here, we exclusively differentially modulated foveal SC state without otherwise differentially altering extrafoveal pre-stimulus SC state (Fig. 2, S4 Figure). Therefore, we now had the perfect opportunity to check how the enhanced visual responses of Figs. 3, 4 looked like when we considered the exact temporal relationship between extrafoveal visual stimulus onset and individual microsaccades (the first two example trials of Fig. 1B show cases in which the stimulus onset happened to occur right before microsaccades towards the extrafoveal stimulus location, demonstrating that we could indeed investigate both pre-and post-microsaccadic modulations).

We measured visual burst strength in visual-motor neurons (like in Figs. 3, 4), but we now related it to when extrafoveal visual stimulus onset occurred relative to individual microsaccade occurrence (Fig. 5; Methods). In the control condition (gray), we replicated our previous observations (31, 34). During foveal retinal image stabilization, we got qualitatively the very same results (blue and red; orthogonal stabilization also gave similar observations), suggesting that the results in Figs. 3, 4 indeed tapped into the same peri-microsaccadic modulations that we had previously documented in the context of covert visual attention shifts (4, 30, 31, 32). Remarkably, foveal retinal image stabilization in the same direction as the upcoming extrafoveal stimulus resulted in two distinct quantitative modifications of the effects: pre-microsaccadic enhancement was strengthened (blue curves; the light blue shows the subset of trials in the same condition in which the actual microsaccade was towards the stimulus, which was the most frequent observation), and post-microsaccadic suppression was significantly reduced (compare blue/light blue and red/gray curves). The former observation is consistent with a directionally-specific effect of pre-movement enhancement (31); the latter is very intriguing because it suggests that the “perpetual” movement plan dictated by foveal retinal image stabilization had a fairly strong alleviating effect on saccadic suppression. The net result was that visual responses were, overall, stronger in the same trials than in the control or opposite ones (Figs. 3, 4).

**Figure 5.**
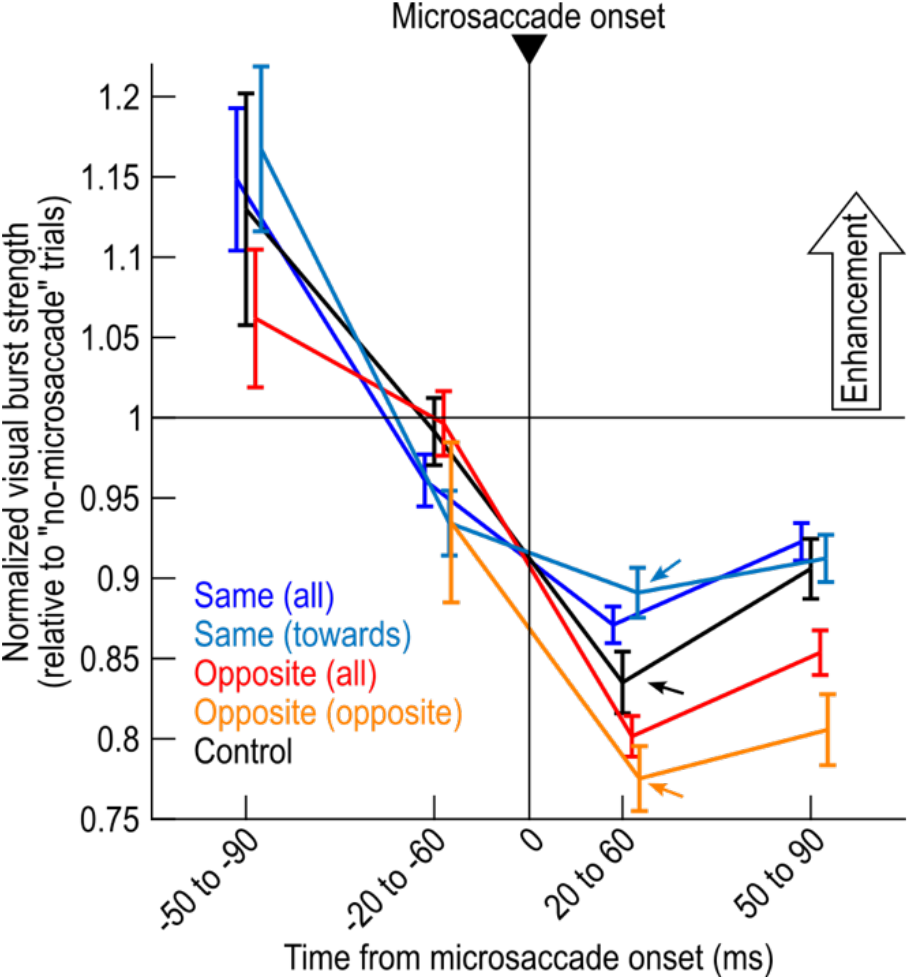
The influence of foveal action on extrafoveal visual sensitivity is mediated by amplifying known peri-microsaccadic changes in extrafoveal SC visual responses. We measured, in visual-motor neurons, visual burst strength (y-axis) as a function of when extrafoveal visual stimulus onset occurred relative to a given microsaccade (x-axis). In the control condition (gray), we replicated prior findings of pre-microsaccadic visual sensitivity enhancement in extrafoveal SC neurons (31, 34). Foveal retinal image stabilization amplified these effects in two ways: before microsaccades, visual sensitivity enhancement (31, 34) was stronger in the same (blue) than opposite (red) condition, and particularly when the microsaccade directions were indeed congruent with the appearing extrafoveal visual stimulus (light blue); after microsaccades, suppression of visual sensitivity due to the microsaccades (31, 80, 81) was substantially weaker in the same than opposite conditions, especially when only the genuine opposite movements in the opposite condition were included (orange; like the approach we took in Fig. 2D). In both cases, this gave rise to stronger visual bursts for same than opposite trials on average (Figs. 3, 4). Error bars denote SEM (Methods). If a time bin has no symbol for one of the curves, it means that there were no data points within that time bin for that specific curve. The underlying data are included in S1 Data.

Statistically, we confirmed the above observations. We ran a two-way ANOVA on the data of Fig. 5 with the factors being: time of stimulus onset relative to a microsaccade and foveal retinal image stabilization condition (control, same, opposite, or same with only the towards movements). We found significant main effects of time (df=3; F=57.79; p=0) and retinal image stabilization condition (df=3; F=2.68; p=0.0455), as well as significant interactions (df=9; F=2.57; p=0.006). For completeness, we also added to Fig. 5 the data in which there were genuine opposite microsaccades in the opposite condition (orange curve). This allowed us to confirm that the slightly increased variance in microsaccade directions in this condition (S3C, D Figure) was not altering our overall results.

Therefore, our results so far confirm that foveal oculomotor state (Figs. 1, 2) is sufficient to differentially modulated extrafoveal visual sensitivity (Figs. 3, 4). Such modulation, remarkably, happens in the same way through which normal microsaccades (without any retinal image stabilization) might be associated with extrafoveal visual sensitivity. This is also consistent with how microsaccades influenced the foveal SC state in the same way independently of retinal image stabilization (Fig. 2D). That is, our retinal image stabilization observations (Figs. 3, 4) were mediated via known peri-microsaccadic changes in visual sensitivity (Fig. 5). These results suggest that the enhnancements seen in Figs. 3, 4 are not fully explained by arousal or target expectation alone, since we observed suppressed, rather than enhanced, visual sensitivity when the stimuli appeared after microsaccades (Fig. 5). More importantly, these results also suggest that at least one component of our previously described links between individual microsaccades and covert visual attentional modulations (1, 4, 30, 32) may indeed be an outcome of how the mere act of microsaccade generation can alter extrafoveal visual sensitivity (4, 31): foveal action is sufficient to differentially modify extrafoveal visual sensitivity.

### Dependence of extrafoveal contrast sensitivity on voluntary microsaccades

If the above hypothesis is reasonable, then we might expect pre-microsaccadic enhancement of human extrafoveal contrast sensitivity in non-attentional tasks, just by virtue of microsaccade generation (4, 30). To test this here, we used a paradigm in which covert visual attention could not be proactively allocated at the tested extrafoveal stimulus location. On every trial, human participants fixated a small white fixation spot (Methods). Then, two identical gabor gratings appeared at 7 deg eccentricity either to the right or left of fixation (82). After a short time, the fixation spot was jumped to the right or left by only ∼10 min arc relative to instantaneous eye position (Methods), thus allowing the participants to voluntarily (57) generate a microsaccade. This aspect of the task was simply like any visually-guided saccade task, except within the foveal scale. At a random time, a test grating appeared on only one side, and subjects evaluated whether it had higher or lower contrast than the standard gratings (Fig. 6A). On control trials, we did not jump the fixation spot; the microsaccades were thus not experimentally controlled, but we excluded trials with peri-stimulus movements because the results of Fig. 5 revealed that they would, in principle (also see below for actual evidence), lead to similar modulations (Methods).

**Figure 6.**
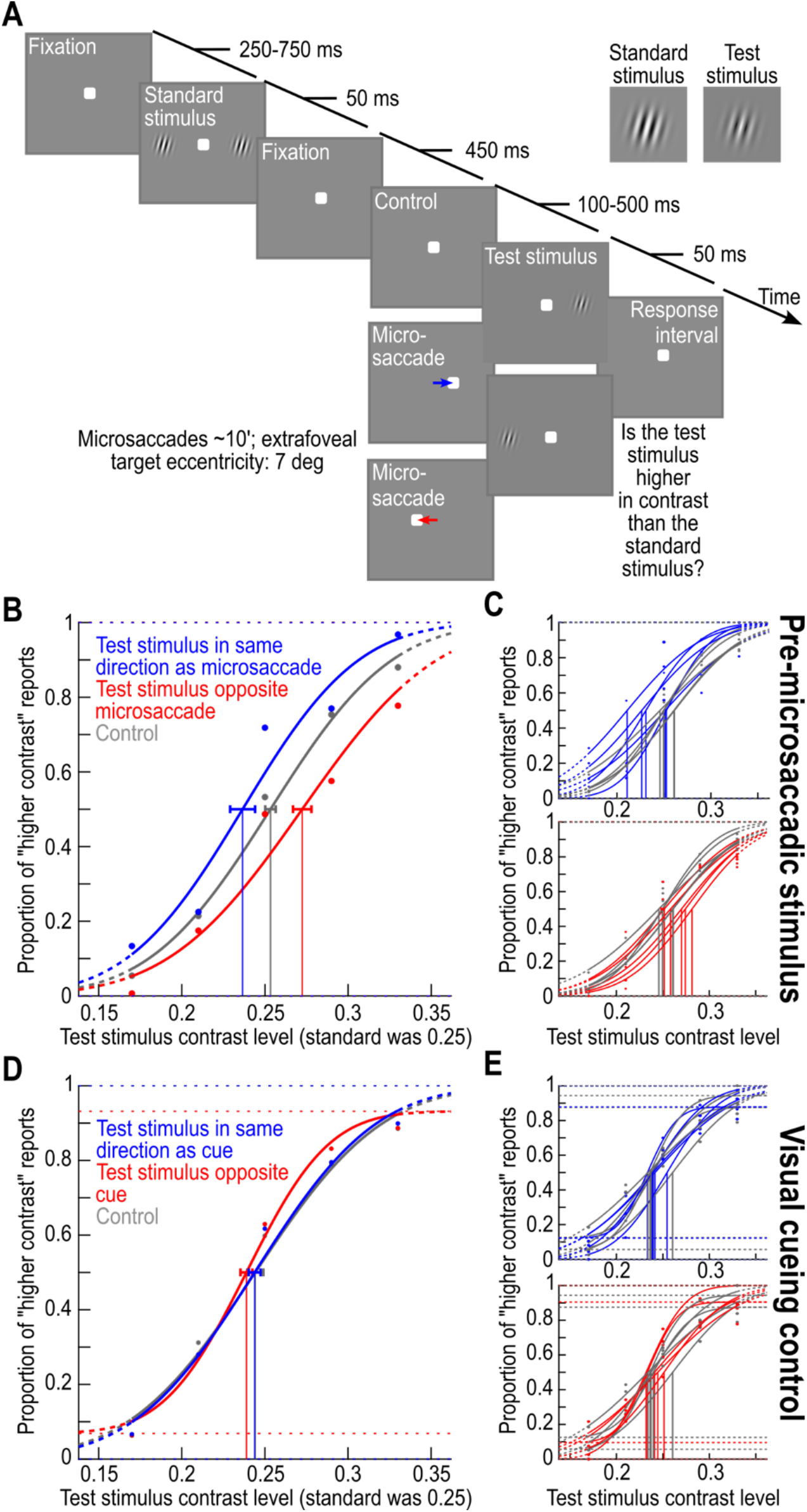
Human voluntary microsaccade generation differentially modulates extrafoveal visual contrast sensitivity in the absence of cued covert attention shifts. **(A)** Humans fixated a central spot. They were then shown a brief, extrafoveal standard stimulus on either side of fixation, so that they could not predictively allocate their attention to either side of the display. After some delay, other than in the control trials, the fixation spot was jumped by a tiny amount either to the right or left, and the subjects were instructed to keep their gaze on it. At a random time, a brief test stimulus appeared extrafoveally, this time on only one side of the display. Subjects assessed whether the test stimulus was higher or lower in contrast than the standard stimuli. **(B)** For test stimuli appearing within 50 ms before microsaccade onset in the same direction (blue), contrast sensitivity was higher than in control. If the test stimuli were in the opposite direction from the microsaccade (red), contrast sensitivity was lower than in control. Error bars denote SEM, and the curves are the average curves across all subjects. **(C)** Individual subject psychometric curves from **B**. The effects of microsaccade direction relative to extrafoveal stimulus direction were highly robust across individuals. **(D)** In a control experiment, the fixation spot remained in its place, and only a transient flash was presented to the right or left of it (identical in location to the fixation spot jump in the main experiment of **A**) (Methods). Contrast sensitivity was unaffected by the foveal visual transient. Thus, the visual transient associated with the fixation spot jump in **A** could not explain the results in **B**, **C**. **(E)** Individual subject results from the experiment in **D**. S10 Figure shows individual subject reaction times and post-microsaccadic stimulus results; S11 Figure shows how similar observations could be made in the control condition, along the same lines of Figs. 2, 5 with the neuronal analyses.

One advantage of this task is that it allowed us to take the primary conclusion from our neurophysiology, namely the role of peri-microsaccadic modulations (Fig. 5), and test it at the perceptual level in humans by looking for peri-microsaccadic contrast sensitivity changes. One other important advantage of the task is that we could now eliminate the arousal and target expectation components of our neurophysiological experiments (e.g. S3, S6 Figures), which were an unavoidable side effect of our retinal image stabilization manipulation (as well as the control of stimulus location per session).

All subjects successfully generated the instructed microsaccades (S10A Figure). Critically, when the extrafoveal test grating appeared within <50 ms before microsaccade onset, contrast sensitivity was enhanced (relative to control) if the grating was congruent with the upcoming microsaccade direction (Fig. 6B, blue), and it was suppressed if the grating appeared opposite the microsaccade direction (Fig. 6B, red) (congruent versus control: p=0.0023; opposite versus control: p=0.0003; permutation test with 10,000 permutations; Methods). This differential modulation of extrafoveal contrast sensitivity was specific to the thresholds, because the width parameters of the psychometric curves were similar across all conditions (control: 0.1881, [0.1397, 0.1916] 95% confidence; congruent: 0.1796, [0.1127, 0.2004] 95% confidence; opposite: 0.2082, [0.1674, 0.2506] 95% confidence). This differential modulation of extrafoveal contrast sensitivity, which was highly robust across participants (Fig. 6C), also did not occur when the test grating appeared within <50 ms after microsaccade onset (S10B, C Figure). Thus, consistent with the impacts of foveal action on extrafoveal visual sensitivity in monkeys (Figs. 1–5), and consistent with our previous results (30, 32), the mere act of generating microsaccades in our human subjects was sufficient to differentially modulated extrafoveal contrast sensitivity (even without a priori attentional allocation).

We also considered the possibility that the fixation spot jumps in the above paradigm (Fig. 6A) might have acted as some sort of visual cue (albeit foveal). While it is not clear how such a foveal transient could alter contrast sensitivity at an eccentricity ∼41 times larger in magnitude than its eccentricity, and specifically only pre-microsaccadically, we next conducted a control experiment in which the participants always maintained fixation on the central spot, and we transiently flashed a small visual spot at ∼10 min arc eccentricity (Methods). This way, we still had a foveal visual transient, but no systematic microsaccades towards it (since the participants were asked to ignore it); we also explicitly excluded all trials in which a microsaccade may have occurred near any stimulus onset in the trials, whether cue or standard/test stimuli (Methods). Moreover, results like those in S3A Figure would suggest that there should be a reduced likelihood of microsaccades after the cue (44). Contrast sensitivity was not altered by the direction of the foveal visual transient (Fig. 6D, E) (congruent versus control: p=0.9472; opposite versus control: p=0.3177; permutation test with 10,000 permutations). Similarly, there were no systematic effects on the widths of the psychometric curves (control: 0.1969, [0.1253, 0.2095] 95% confidence; same: 0.1888, [0.1176, 0.1987] 95% confidence; opposite: 0.1283, [0.1030, 0.187] 95% confidence). Thus, the effects of microsaccade generation in Fig. 6B, C were not due to our introduction of foveal visual transients by the fixation spot jumps.

Finally, just like we did with our neuronal analyses, we wondered whether congruent microsaccades during the control condition could still be associated with similar pre-microsaccadic modulations to those in Fig. 6B, C (replicating our earlier findings (30, 31, 34)). That is, even though microsaccades in the control trials were not experimentally controlled, some of them were necessarily either congruent or incongruent with extrafoveal visual stimulus location, and with the proper timing relationship to the stimulus onset. When we analyzed these microsaccades, we indeed found that similar results to those in Fig. 6B, C could be possible (S11 Figure). Therefore, microsaccades are indeed associated with long-ranging effects (in terms of eccentricity) on extrafoveal contrast sensitivity, even in the absence of any systematic covert attentional allocation.

### Spread of contrast sensitivity enhancement ahead of larger saccade endpoints

If microsaccade generation is sufficient to modulate extrafoveal visual sensitivity ahead of the movement endpoints, then one might predict that contrast sensitivity can also be differentially modulated by larger saccades, and ahead of the saccade endpoint locations (i.e. at larger eccentricities). To test this, we designed a second human task similar to that in Fig. 6, but this time instructing larger 5-deg saccades. We found that pre-saccadic contrast sensitivity was enhanced ahead of the saccade target for congruent than incongruent saccades, even when controlling for the retinal positions of the pre-saccadic test stimuli (Methods; S12 Figure). These results replicate prior mappings of pre-saccadic contrast sensitivity (83, 84, 85), and, for our present context, they demonstrate that it is indeed possible for the process of saccade and microsaccade generation to influence visual sensitivity even at eccentricities ahead of the eye movement endpoints. This supports our interpretation of a plausible causal influence of foveal action on extrafoveal vision.

## Discussion

We found that exclusive experimental modulation of foveal oculomotor state was sufficient to alter extrafoveal visual sensitivity and behavior (Figs. 1–4, 6). Moreover, this alteration was mediated by known peri-microsaccadic changes in visual sensitivity (Figs. 5, 6) (31, 34).

Our monkey paradigm held the monkeys’ expectational state about the extrafoveal stimulus location constant. This is not different from the monkey studies (11, 12) that debated the recent Lowet et al. (13) claims about microsaccades and attention. Indeed, in these monkey studies (11, 12), a “cue” was redundant because the experimental conditions were blocked, raising the question of why such a cue was used in the first place. We think that, in these experiments, there was an altered internal state due to the blocked task design, not unlike the evidence that we saw in our own experiments, whether in the microsaccades (S3 Figure) or in the saccadic reaction times (Fig. 4E, S6 Figure). Regardless of whether this can be called attention or not (86), it represents a large departure from the original description of microsaccades as tracking covert attention “shifts” (1). In a very similar manner to these studies, we also had evidence of an altered internal state during the pre-stimulus epoch (S4D-F Figure). However, this altered state was not differentially modulated by foveal oculomotor state, and it was therefore not adequate to explain our differential sensitivity effects (Figs. 3, 4). It was also likely due to arousal (S3A Figure), target location knowledge (S6 Figure), and the temporal cue of retinal image stabilization onset (S3B-D Figure). Thus, even though it is described in these studies (11, 12) as the neuronal correlate of attention, we view such altered steady-state activity in these experiments as being orthogonal to the debate regarding microsaccades and attention “shifts”.

Concerning the Lowet et al. (13) findings, we suggest that they reflected the simple fact that microsaccades are, at the end of the day, eye movements. In their experiments, the cue was a punctate line near the fovea. Our behavioral results (Fig. 1), and others from the literature (40), suggest that this line is a very potent stimulus to guide microsaccades towards its endpoint. Hence, it is not too surprising that their microsaccade directions were highly congruent with the cue direction; the microsaccades were simply “foveating” eye movements. Critically, by virtue of being eye movements, these microsaccades brought the extrafoveal stimuli closer to the fovea. Given that visual receptive fields are skewed (having longer tails with increasing eccentricity) due anatomical retinotopic map distortions (52, 87), shifting stimuli slightly closer to the fovea can elevate population neuronal firing rates. This would constitute a simple visual mechanism that, in our opinion, can parsimoniously explain results like those of Lowet et al. (13). It is quite distinct from our peri-movement effects.

More broadly, acknowledging that microsaccades are genuine eye movements can also make sense of potential “deviations” from their expected (and known) relationships to attention shifts. Such deviations have led to suggestions that microsaccades are only “functionally” but not “obligatorily” (12, 16, 18) linked to attention. However, by the very same logic, even large saccades should be described as only having a “functional” but not “obligatory” link to attention, despite clear evidence that these eye movements causally alter vision, whether through peri-saccadic changes in representations (88), or through pre-saccadic attention (82, 83, 89), or even simply through the allocation of expanded foveal resources in magnified retinotopic maps. For example, large saccades robustly land in between two different stimuli (but not on either of them individually) (90), just like microsaccades do (8, 11). This does not, in any way, mean that large saccades are suddenly no longer linked to attention or to the act of foveation. Instead, it means that saccade endpoints reflect the overall landscape of activity in the SC (and related structures) (45). Indeed, it can be said (without much controversy) that large saccades both reflect visual, motor, and cognitive guidance as well as influence the processing of visual, motor, and cognitive signals. We argue that this same sentence is also true for microsaccades. In fact, there is now enough neurobiological and behavioral knowledge to allow predicting (almost deterministically in some cases) when and to which direction microsaccades would happen in a variety of sensory and cognitive tasks (4, 10, 32, 33, 44, 46, 47). Coupled with known peri-microsaccadic changes in sensitivity like those in Fig. 5, and in other forms (42), this means that it may indeed be technically possible to predict (or “track”) visual perceptual performance changes in tasks by monitoring microsaccades. We believe that this sentiment is not controversial, and it is also pragmatic: it favors circuit mechanisms over debates about what may or may not be called attention. Most importantly, one can additionally use foveal retinal image stabilization, like we did here, to even explicitly manipulate extrafoveal visual sensitivity.

From a more fundamental perspective, we find it intriguing that extrafoveal eccentricities can be affected by much smaller eye movements. We think that wide-field SC cells (91) could be one way to mediate such an influence. It would be interesting to explore this further, especially in the primate SC. Indeed, we recently identified a reverse transfer of visual information from the extrafoveal SC representation to the foveal one across large saccades (92), and it has also long been known that there are traveling wavefronts in the SC at the time of large saccades (93). Even more recently, we found evidence that foveal SC transients seem to be critical for jumpstarting extrafoveal saccadic orienting (94). Thus, there are precedents to an impact of one locus in the SC on other, distant, loci, and also ultimately on perceptual performance (by virtue of this structure’s important role in dictating behavior).

We are also fascinated by the observation that foveal retinal image stabilization did not modulate visual-only SC neurons (S7 Figure). In our previous work, we observed pre-microsaccadic enhancement for both visual-only and visual-motor neurons, although with slightly different properties (31). Our current results suggest that colliculo-collicular effects like we emphasized in our present experimental design (Fig. 2) may be restricted to only the visual-motor SC layers. Thus, modulations of visual-only neurons could be mediated by external inputs to the SC.

Finally, we found that post-microsaccadic suppression was significantly alleviated by our “perpetual” microsaccade command (Fig. 5). Saccadic suppression is, in general, difficult to weaken; once it happens, it happens strongly, likely owing to its visual origin (95, 96, 97). Thus, observing its alleviation with a forced foveal visual error is an interesting finding, and it could be exploited in tasks in which it would be desirable to shorten, or otherwise lessen, the disruptive effects of saccadic suppression.

## Methods

### Ethics statement

All animal experiments were approved by the local governmental authorities of the city of Tübingen (Regierungspräsidium Tübingen), under licenses CIN3/13 and CIN4/19G. The experiments complied with the European Union directives on animal research, as well as the national implementations of these directives.

All human experiments were approved by the ethical committee for human experimentation at the Faculty of Medicine of the University of Tübingen. The participants were compensated financially for their time, and they provided written, informed consent. The experiments were in accordance with the Declaration of Helsinki.

### Experimental animals

We collected behavioral data from four adult, male rhesus macaque monkeys (A, aged 9 years; F, aged 8 years; M, aged 9 years; and N, aged 10 years). We also collected neuronal data from three of these monkeys (A, N, and M) using either single electrodes (monkey M) or linear multi-electrode arrays (monkeys A and N). All three monkeys were shown to exhibit pre-microsaccadic sensitivity enhancements in other experimental contexts (N: Chen et al., 2015 (31); A and M: Wu and Hafed, 2025 (34)), increasing our confidence in the present results.

The animals were all previously prepared for behavioral and neurophysiological experiments. Briefly, each animal underwent surgeries, under full anesthesia, to receive head-holder implants, chamber implants (and craniotomies), and scleral search coils. The recording chambers were placed centered on the midline of the skull and tilted backward from vertical by an angle of 38 deg. The chamber centers were aimed, based on structural MRI images, to allow electrodes a straight path towards the superior colliculus (SC), and we could target both the right and left SC in each animal using the same midline chamber. The scleral search coils allowed us to measure eye movements with very high precision using the magnetic induction technique (98, 99).

### Laboratory setup for animal experiments

The experiments for monkey N were conducted using the control system described in refs. (8, 32, 56). Briefly, we used a real-time computer from National Instruments (cRIO-9024), paired to high-speed digital I/O and analog-to-digital converter cards. The system controlled all aspects of data acquisition and reward delivery, and it also commanded a graphics computer using the PsychToolbox (100, 101, 102) for displaying images to the monkey. The control system ran at 1KHz sampling rate, and the display monitor (CRT) had a refresh rate of 120 Hz. Data was stored using the Plexon Multi-acquisition processor (MAP) device.

For the experiments in the remaining monkeys, we used a custom-built modification to PLDAPS (103), which again used the PsychToolbox for display control. Data were stored using the OmniPlex system from Plexon, and the display refresh rate was either 120 Hz or 85 Hz.

For real-time retinal image stabilization, we used the methods mentioned earlier (32). Specifically, we calibrated eye position by having the monkey fixate 19 different locations on the display multiple times. We then obtained multi-order polynomials converting the raw measurements (arriving at 1 KHz rate) into calibrated eye positions. Before the start of retinal image stabilization on every trial, we always performed a final offset correction by measuring the final 50 ms of eye position (and averaging it) before initiating retinal image stabilization. This allowed high-fidelity retinal image stabilization during gaze fixation. We successfully used this system before, to control foveal errors (32, 33) like we did here, and also to control extrafoveal stimulus retinal image positions (48).

In all experiments, the monkeys sat in a dark room, and the CRT display in front of them had a gray background (21 cd/m^2^ for monkey N and 26.11 cd/m^2^ for the others). Stimuli like the fixation spot and the extrafoveal visual target were white (86 cd/m^2^ for monkey N and 79.9 cd/m^2^ for the others). Also, the distance to the display was 45 cm in monkey N and 72 cm in the other monkeys, resulting in display pixel resolutions of ∼22 pixels/deg for monkey N and ∼34 pixels/deg for the other monkeys.

Finally, electrodes consisted of tungsten microelectrodes (from FHC) in the case of single-electrode recordings (1-1.5 MOhm impedances), and they were 16- or 24-channel V-Probes (from Plexon) in the case of linear electrode array recordings.

### Experimental procedures with the animals

The behavioral task for the monkeys consisted of a modified visually-guided saccade task. Each trial began with a fixation period on a white, central fixation spot. The fixation spot was a square of ∼10.8 x 10.8 min arc dimensions in monkeys A, F, and M, and it was a square of ∼8.5 x 8.5 min arc dimensions in monkey N. After fixating the spot for a random interval between 450 ms and 650 ms, we first sampled eye data for 50 ms (at 1 KHz sampling rate) to prepare for offset correction in the retinal image stabilization manipulation. Then, on non-control trials, we used real-time retinal image stabilization to update the position of the fixation spot on the display with every single frame refresh. Such updating was contingent on the instantaneous eye position, such that we always displayed the fixation spot with a constant offset relative to the measured line of sight. This constant offset created a “perpetual” foveal visual error, which was effective in driving microsaccades attempting to minimize this error. After 150-500 ms, we removed the fixation spot and displayed an extrafoveal stimulus, consisting of a white disc with radius 0.51 deg. This extrafoveal stimulus was to be immediately foveated by the monkeys, and the monkeys needed to hold fixation on it for at least 500 ms in order to receive the final reward.

The extrafoveal stimulus appeared at a single retinotopic location, which was dictated by the recorded neurons’ response field (RF) locations, within any given session (on sessions with behavior only, we picked a location for the extrafoveal visual stimulus in one of the four visual quadrants for each session). Note that we also took care to account for instantaneous eye position at the time of extrafoveal visual stimulus onset. That is, if the eye was deviated slightly from display center, then the extrafoveal stimulus was also deviated by the same amount (but no further retinal image stabilization after the stimulus onset was applied). This was very important because if we had only picked a single stimulus location in display coordinates (irrespective of eye position), then our stimulus onset would have been at different locations relative to neurons’ RF’s, which would have, in turn, reduced and blurred our visual responses (48, 104). Moreover, a deviation of eye position during retinal image stabilization is inevitably expected when such retinal image stabilization involves the fixation spot. This is because the oculomotor system attempts to continuously, but unsuccessfully, eliminate the imposed forced foveal visual error (S1 Movie shows an extreme example of this if a large visual error is imposed during retinal image stabilization).

When we were recording foveal neurons, we placed the extrafoveal stimulus at a position between 3.54 and 12.81 deg eccentricity (dictated by extrafoveal electrode positions during simultaneous recordings), and we chose its direction to coincide with the directions of the foveal neurons that we were recording from. This way, a forced foveal visual error in the same direction as the extrafoveal stimulus was at the same time a foveal visual error that should have activated the foveal neurons if microsaccades were triggered in their direction. When we were recording extrafoveal neurons, the extrafoveal stimulus location was at the location where most RF’s in the recording session were encountered. We assessed the RF’s online during the sessions, using standard visually-guided and memory-guided saccades (31, 70).

On the retinal image stabilization trials, we had four different conditions. In one condition, the forced visual error direction was in the same direction as the upcoming extrafoveal stimulus. In another, it was diametrically opposite from it. And, in the remaining two conditions, it was a direction that was rotated by 90 deg clockwise or counterclockwise from the direction of the extrafoveal visual stimulus. Of course, because displays are pixelated, we had to bin the extrafoveal stimulus directions when calculating the pixel offset. S1 Figure shows the details in such binning. For example, if the extrafoveal stimulus was within +/- 11.25 deg in direction from the rightward horizontal axis, then the pixel offset that we applied relative to instantaneous eye position was 2 pixels to the right of the current eye position. If the extrafoveal stimulus was within +/- 11.25 deg from a direction of 22.5 deg above the rightward horizontal axis, then the pixel offset that we applied was now 2 pixels to the right and 1 pixel above the current instantaneous eye position, and similarly for other RF directions. This way, we had sufficient directional resolution in our pixel offsets to allow recording from all visual quadrants. Thus, in monkey N, the fixation spot offsets (in visual angles) were in the range of 5.5-7.7 min arc, and for the other monkeys, they were in the range of 3.5-5 min arc.

For Monkey A, we completed 44 sessions with a total of 23,463 trials. Twelve of the sessions included neuronal recordings; the rest were used as additional data for our behavioral analyses. For Monkey M, we conducted 32 sessions with a total of 30,861 trials. 23 of these sessions used single-electrode recordings. For Monkey N, we completed 27 sessions with 14,713 trials, all of which included neuronal recordings. In this monkey, only trials from the control condition were previously analyzed for a different scientific context (45). Finally, we also collected behavioral data from Monkey F across 9 sessions with 15,595 trials.

### Data analysis for the animal experiments

We detected all microsaccades and saccades using our previously established methods (56, 105).

To assess the success of retinal image stabilization, we checked the distribution of microsaccade directions in each forced visual error condition. To do so, we calculated microsaccade direction relative to the direction of the extrafoveal visual stimulus, and we then plotted directional histograms (with angular bins of 12 deg size). We then compared these directional histograms to those obtained from the control condition. We also characterized the variance in microsaccade directions in the different conditions (S3C, D Figure). For each axis in which there was a clear direction bias (e.g. towards the extrafoveal stimulus in the same condition or 180 deg from it in the opposite condition), we plotted a normalized histogram of microsaccade directions relative to this axis. This gave a peaked distribution when retinal image stabilization was successful. From this peak distribution, we calculated the standard deviation across all microsaccades. To show that there was a difference in the standard deviation of microsaccade directions across conditions (S3C Figure), we bootstrapped the measurements (10000 times) to obtain 95% confidence intervals on the estimated standard deviation measurements (S3D Figure).

We also checked the time course of microsaccadic modulations after the onset of retinal image stabilization. We divided all microsaccades into four different categories depending on their direction relative to the upcoming extrafoveal stimulus location. That is, if the microsaccade had an angle within +/-45 deg from the stimulus, it was classified as same; if it had an angle within +/- 45 deg from the opposite direction of the stimulus, it was classified as opposite; and so on. Then, we plotted the fraction of microsaccades within a given category as a function of time from retinal image stabilization onset. If the microsaccades were equally likely to any direction before retinal image stabilization, then this time course should have been hovering at a fraction of around 0.25. We used a binning window of 50 ms, which we moved in steps of 5 ms, to generate these time courses.

Finally, we also analyzed microsaccade amplitudes and plotted them across conditions, both individually for each monkey (S2 Figure) and also across all monkeys (Fig. 1E, F), and we calculated microsaccade rates (S3 Figure A, B). To do so, we used similar temporal binning as we did for the microsaccade direction time courses (50 ms time windows, in steps of 5 ms).

For the large saccades towards the extrafoveal visual stimulus, we measured their reaction times as a function of the different experimental conditions. We also characterized the likelihood of express saccades. We defined those as the movements with reaction times shorter than 100 ms. In analyses in which we related neuronal sensitivity to reaction times, we included all trials (including those with express reaction times). In terms of landing positions, we only accepted trials in which the saccades landed within 2 deg from the extrafoveal stimulus location.

Neuronally, we sorted units online during single electrode recordings. For the rest, we sorted the neurons using KiloSort (106).

For our extrafoveal neuron database, we only included neurons that exhibited a visual response. We assessed such a response as follows. We defined two intervals: a pre-stimulus interval consisting of the final 100 ms before extrafoveal stimulus onset on control trials, and a post-stimulus interval from 50 ms to 100 ms after stimulus onset. If average firing rate during the post-stimulus onset was significantly elevated relative to the average firing rate during the pre-stimulus onset across trials (using a paired t-test), we included the neuron in our database.

For our foveal neuron database, we included neurons with foveal RF’s and microsaccade-related discharge (24). These neurons did not possess visual responses to the extrafoveal stimulus onset.

We had two classes of neuronal analyses, one for pre-stimulus activity and one for stimulus-evoked visual responses. First, we analyzed pre-stimulus activity, to investigate the impacts of retinal image stabilization on both the foveal and extrafoveal SC state. For the foveal neurons, we plotted saccade-free activity in the final 300 ms before extrafoveal stimulus onset. By saccade-free, we mean that we excluded any neuronal activity within 50 ms on either end of a microsaccade. To summarize the effects across the population, we first normalized the activity of each neuron before averaging across neurons. The normalization factor was the average saccade-free foveal neuron activity in the same condition (because it generally had the highest activity out of all conditions). To obtain scatter plots of raw firing rates, we averaged firing rate in each neuron during the interval from -150 ms to -50 ms relative to extrafoveal stimulus onset. To analyze microsaccade-related activity in the foveal neurons, we collected all microsaccades occurring in the final 300 ms before extrafoveal stimulus onset in the control condition. In retinal image stabilization conditions, we included all microsaccades occurring between the onset and end of foveal retinal image stabilization. For some analyses, we included all microsaccades in a given condition (e.g. same or opposite), regardless of the microsaccade directions in the condition. If the manipulation of foveal retinal image stabilization was successful, then expected neuronal modulations were to be observed. For example, same trials would exhibit microsaccade-related motor bursts since the manipulation increased the likelihood of microsaccades in the same direction as the RF. However, to better control for microsaccade directions, especially for different ranges of direction variances across the different conditions (S3C, D Figure), we also constrained microsaccade direction ranges in some analyses (to within +/- 45 deg from a given axis). This allowed us to have better estimates of the observed neuronal modulations, without potential masking by unrelated microsaccade directions. We also followed a similar strategy in extrafoveal neuronal analyses as well.

For the pre-stimulus analyses of extrafoveal neurons, we again measured saccade-free neuronal activity in the final 300 ms before extrafoveal stimulus onset. We also applied a similar normalization factor to average pre-stimulus activity across neurons. However, for these neurons, the normalization factor was always the peak, baseline-subtracted stimulus-evoked visual burst magnitude in the control condition (we did this because we used this same normalization factor in our main analyses of extrafoveal visual burst strengths described below). Also, for scatter plots of raw firing rates in the pre-stimulus interval, we again used the same measurement interval as just described above. Note that the use of peak visual response is standard practice. Also, given that we collected many trials per condition (e.g. Fig. 3), and given that we explicitly tested our neurons for having a significant visual response (above), then taking the peak of an across-trial average firing rate curve is not prone to too much noise and variability.

We next analyzed post-stimulus activity in the extrafoveal neurons, focusing on our main goal of assessing the strengths of stimulus evoked visual bursts. Here, we first calculated baseline-subtracted firing rates, where the baseline was calculated as the average firing rate over the 50 ms interval before extrafoveal stimulus onset. After subtracting the baseline, we normalized all data points by dividing them by the peak visual response in the control condition, defined as the maximum firing rate within 80 ms of extrafoveal stimulus onset. Scatter plots measured the peak visual response in the interval from extrafoveal stimulus onset to 80 ms. We also calculated neuronal modulation indices (for example, comparing the same to control conditions). These indices were calculated as the peak visual burst for one condition (same) minus that for the other condition (control), divided by the sum. If the indices were positive, then there was enhancement in visual sensitivity relative to control; if they were negative, then visual sensitivity was suppressed. In all analyses, whenever we measured visual responses in the control condition, we always excluded trials in which there were microsaccades during the interval from -50 ms to +30 ms relative to extrafoveal stimulus onset. We also included individual monkey neuronal results.

To relate visual burst strength to the time between stimulus onset and microsaccade onset (31, 34), we used an approach similar to that we employed previously (34, 80), in the sense that we normalized each neuron’s response and then pooled all responses across neurons. This allowed us a more robust estimate of peri-microsaccadic modulation time course despite limited numbers of trials. From the control condition, we took all trials in which there were no microsaccades within 300 ms from extrafoveal stimulus onset. We then measured peak firing rate as above, and used it as the normalization factor. Then, for each condition (including control) in which there were microsaccades within a specific time window relative to stimulus onset, we again measured the peak firing rate and divided it by the normalization factor. This allowed us to observe the conditions under which visual burst strengths were either enhanced or suppressed around microsaccade onset.

We had a total of 400 extrafoveal neurons (361 visual-motor and 39 visual-only; classified as we had done previously (81)) and 41 foveal neurons in our database.

### Human participants

For the voluntary microsaccade generation experiment (e.g. Fig. 6), we recruited six human participants (five females and one male), aged 22-26 years. All, but author TZ, were naïve to the purposes of the experiment, and they all gave written and informed consent. Three subjects were experienced with psychophysical experiments, while the rest were participating for the first time. We excluded the male subject from the study because he was unable to successfully perform the comparison between standard and test stimuli in the task (task details are provided below and in Fig. 6). For the rest of the participants, each of them completed six sessions (on different days) of the main experiment and two sessions of the control experiment (both described in more detail below), with each session lasting between 45 and 60 minutes each.

For the large saccade experiment (S12 Figure), we tested eight subjects (2 male and 6 female), aged 24-45 years. Two of the subjects had participated in the voluntary microsaccade generation experiment earlier.

All subjects provided written and informed consent before participating in this experiment, and they were also financially compensated for their time. Each subject participated in 10-11 sessions for this experiment.

### Laboratory setup for human experiments

We used the same laboratory environment as that used in our recent studies (107). Briefly, the subjects sat 57 cm in front of a calibrated CRT monitor having 85 Hz refresh rate and ∼41 pixels/deg resolution. We used a video-based, desk-mounted eye tracker (EyeLink1000; SR Research, Inc.) having a sampling rate of 1KHz, and the room was otherwise dark. To improve eye tracking quality, we fixed the subjects’ head position using a custom-built apparatus with mounts for stabilizing the position of the chin, forehead, temples, and back of the head (30). The subjects responded by pressing a button on a hand-held response box.

Stimuli were presented on a gray background of luminance 22.15 cd/m^2^. The fixation spot was a small, white (95 cd/m^2^) square spanning ∼4.4 by 4.4 min arc. A similar spot served as the eye movement target when microsaccades/saccades were instructed, and also as the ignored foveal cue in the control version of the microsaccade experiment.

### Experimental procedures with the humans

For the voluntary microsaccade generation experiment, each trial started with a white fixation spot at the center of the display. After 250-750 ms, two identical standard stimuli (gabor grating patches with 25% contrast, 2 cycles/deg frequency, and 0.33 deg σ parameter) were briefly presented for 50 ms and at a 7 deg to the left and right of the fixation spot. The gratings had identical orientation and phase, which were randomized across trials. The subjects were instructed to maintain fixation on the spot, but to note the contrast of the extrafoveal standard stimuli, and they were also told that the stimuli were identical to each other in all respects (similar, in principle, to the experimental design of ref. (82)). After another 450 ms, on non-control trials, we jumped the fixation spot by ∼10 min arc to the right or left of instantaneous eye position. This way, we could maximize the likelihood that a microsaccade was successfully triggered, since we were generating a foveal visual error exactly like we did in the monkey experiments. We estimated instantaneous eye position by taking the median of the eye position samples collected from the eye tracker over the last 50 display frame updates before the fixation spot jump. This way, we had a robust estimate of instantaneous eye position that was more immune to eye tracker noise. The subjects were instructed to follow the fixation spot, just like in any other visually-guided saccade task. On control trials, the fixation spot was not moved at all. After another interval of 100-500 ms, a test stimulus (gabor patch) was briefly flashed for 50 ms at 7 deg either to the left or right of the jumped fixation spot position. The test stimulus was identical to the standard stimuli of the trial, except that it could have one of five different contrast levels (17%, 21%, 25%, 29%, or 33%). After the test stimulus flash, the subjects reported whether they perceived the test stimulus as having a higher or lower contrast than the standard stimuli (they pressed one button for higher and another button for lower). Thus, this was a “forced-choice” experiment, meaning that even if the contrast appeared the same, subjects had to choose either “higher” or “lower”. Also, our choice of variable timing between fixation spot jump and test stimulus flash was explicit: we wanted to have enough trials with “pre-microsaccadic” test stimulus onset. Thus, we estimated the expected microsaccade reaction times (57), and we picked a time range of test stimulus onset that straddled the distribution of these reaction times. This way, we could probe perceptual contrast sensitivity in the immediate pre- and peri-microsaccadic intervals. The experiment included three conditions: (1) “same,” where the test stimulus appeared in the same direction as the microsaccade (i.e. the fixation spot jump direction); (2) “opposite,” where the test stimulus appeared in the opposite direction from the microsaccade; and (3) “control,” where no fixation spot jump occurred (microsaccades could still occur, but they were not experimentally manipulated by introducing an explicit foveal visual error). In each session, we ran three blocks of 240 trials each. Within each block, subjects took short breaks every 20 trials if they reported feeling fatigued, and these breaks also allowed for frequent recalibration.

To rule out potential cueing effects caused by the foveal visual transient of fixation spot jumping, we conducted a control experiment for the above paradigm. Instead of shifting the fixation spot, we induced a transient foveal cue by flashing a white square (same as the fixation spot) ∼10 min arc to the right or left of the fixation spot, and for ∼35 ms. This transient flash acted as a foveal cue, and subjects were instructed not to make microsaccades towards it. This was very easy for them to do because the transient was so brief, and there was no other stimulus next to the fixation spot except for the mere few frames of the flash. In the data analyses (see below), we excluded any rare trials in which microsaccades occurred around the onset of the foveal cue. Apart from this modification, all other procedures were identical to those of the main experiment above. Each session included three blocks of 200 trials each.

The experiment with larger saccades was very similar to the main voluntary microsaccade experiment above, with one major difference: instead of prompting subjects to make a small microsaccade, we placed the saccade target 5 degrees away from the fixation point. This required the subjects to prepare a 5 deg saccade before the test stimulus appeared. Also, the test stimulus could appear on either side of the initial fixation spot, at 7 deg eccentricity relative to either the initial fixation spot location or its location at the saccade endpoint (S12A Figure). That is, retinotopically, if the test stimulus appeared in the same direction as the saccade and before saccade onset, then it was at either 7 deg or 12 deg eccentricity (i.e. 2 deg or 7 deg more eccentric than the saccade target location). If it appeared in the opposite direction, then it was at either 2 deg or 7 deg in the opposite direction from the upcoming saccade. The 2 deg case was the result of the 7 deg position relative to the endpoint of the saccade target, and we included it to keep the experimental design symmetric; our comparison of interest was the 7 deg position opposite the saccade direction to the 7 deg and 12 deg positions in the same direction of the saccade (and ahead of the saccade target location). Finally, another minor difference from the main microsaccade experiment above is that we slightly altered the distribution of possible test stimulus times. Specifically, to increase the amount of data available for analysis before saccade onset, we shortened the delay between the saccadic target onset and the test stimulus to the range of 50-250 ms. Finally, in each session, we collected 3 blocks of 250 trials each.

### Data analysis for the human experiments

We detected saccades and microsaccades using our previously described methods (56, 105). In the main microsaccade experiment, we filtered out any trials with saccades larger than 3 deg at any time, which could happen due to rare lapses, distractions, blink-related eye movements, and so on. Since subjects were not allowed to directly look at the extrafoveal stimuli throughout the experiment, large saccades were theoretically impossible, and we only included trials with microsaccades smaller than 0.5 deg in amplitude for the final data analyses. Additionally, in the “microsaccade conditions”, we excluded trials that had no (or more than one) microsaccades between 50 ms and 400 ms after fixation spot jump. Trials with microsaccades occurring within 100 ms around the presentation of the standard stimulus were also excluded to prevent the influences of microsaccades on perception (108). So, the logic of the experiment was similar to classic studies of peri-saccadic vision: there is one instructed saccade (fixation spot jump), and we needed to have one saccade in a given trial in order to be able to measure peri-saccadic changes in performance (the one saccade just happened to be small). For our purposes, we were interested in differential modulations with microsaccade direction, which we believe were not dominated by potentially different cognitive state in this task when compared to the transient cueing case.

From the filtered data, we extracted three conditions: “no microsaccade around extrafoveal test stimulus onset”, “microsaccade in the same direction as the extrafoveal test stimulus”, and “microsaccade in the opposite direction from the extrafoveal stimulus”. We considered a microsaccade to be in the same direction as the extrafoveal stimulus if its horizontal component was to the same hemifield; a microsaccade was opposite if its horizontal component was to the opposite hemifield. For each condition, the proportion of higher button presses at each stimulus contrast was calculated, and the probabilities of “higher” choices across different stimulus levels were plotted as a psychometric function. To determine the threshold in each experimental condition, psychometric functions were fitted using the *psignifit* toolbox (version 2.5.6) for Matlab (109, 110, 111). For our data analysis, we selected the cumulative normal sigmoid function for curve fitting and used it to obtain the thresholds. The threshold was defined as the stimulus level corresponding to 50% of higher contrast reports in the perceptual performance. We calculated the threshold for each subject individually, and, for the microsaccade conditions, we calculated the threshold either for stimuli appearing within <50 ms before microsaccade onset or for stimuli appearing within <50 ms after microsaccade onset. In some analyses, we also explicitly looked for trials in the control condition, during which a microsaccade happened to occur within <50 ms before or after microsaccade onset. We then classified the microsaccades according to whether their directions were congruent and incongruent with the extrafoveal test stimulus location, and we calculated the subjects’ thresholds. Even though the data was less numerous than in our experimentally controlled microsaccade conditions, this allowed us to confirm that the same contrast sensitivity changes that we experimentally tested in the microsaccade conditions could still happen for microsaccades in the control condition. For summary plots, we averaged each corresponding data point across subjects (e.g. at each contrast level), and we then used *psignifit* again to obtain a new fit across the population average measurements. We also calculated the SEM of the thresholds across subjects and indicated in the figures showing population psychometric curves. For completeness, we also measured the widths of the psychometric curves, and their 95% confidence intervals, again using *psignifit*.

To check for statistically significant differences in thresholds between the same and control conditions or the opposite and control conditions, we used permutation tests. Specifically, for each contrast level of the test stimulus, we checked (for each subject) how many trials we had in control and how many trials we had in same (or opposite). We also checked how many trials in each condition had a “higher” response by the subject. Then, we created a “resampled” subject’s results, having the very same number of trials per condition but shuffling the behavioral responses from the original data (without replacement). We then obtained the threshold for same (or opposite) in the resampled subject’s data and the threshold for the resampled control data, and we did this across subjects to get a population threshold in each condition. We repeated this shuffling process 10,000 times, each time getting a threshold difference between same (or opposite) and control. If the true threshold difference in the original data deviated from the distribution of the shuffled differences, then we considered our threshold difference to be significant. We used a p-value of less than 0.05 to assess significance in this way.

The psychometric curves and statistical tests for the cueing control variant of the microsaccade experiment were performed in the same way.

For the large saccade psychophysical experiment, we ensured no microsaccades occurring within 100 ms from the standard stimulus onset. This avoided any alterations in contrast sensitivity due to microsaccades. Then, to confirm that the subjects’ saccades fell within a reasonable range around the saccade target, we plotted the distribution of all saccade landing locations in the horizontal and vertical directions. We calculated the median of this distribution, and we drew a 1.5-degree circle around it. Only saccades landing within this circle were included in subsequent analyses. Retinal coordinates of the pre-saccadic test stimulus were calculated by subtracting the mean eye position from the last 25 ms before saccade onset. We calculated thresholds exactly like above, and we defined sensitivity as the inverse of threshold. The same permutation test procedure was used to calculate p-values, as described above.

## Competing interests statement

The authors declare no competing interests.

## Data availability statement

All relevant data are within the paper and its Supporting Information files.

## Acknowledgements

We were funded by the Deutsche Forschungsgemeinschaft (DFG; German Research Foundation) through the Special Priority Programme: “SPP 2411 Sensing LOOPS: cortico-subcortical interactions for adaptive sensing” (project number: HA 6749/11-1). We were also funded by the Werner Reichardt Center for Integrative Neuroscience, through the DFG’s Excellence Cluster initiative (EXC307).

## Supporting figure/movie legends

**S1 Movie. Demonstration of the effects of retinal image stabilization on corrective rapid eye movements, when using a larger amplitude of forced visual error (much larger than we employed in our actual experiments).** This video shows the eye position of one monkey (N) when suddenly displaying the fixation spot gaze-contingently, at a large forced visual error of ∼1.5 deg either to the right or left of instantaneous eye position. Note how a series of staircase saccades are triggered, in an unsuccessful attempt by the oculomotor system to eliminate the experimentally-induced visual error. This instability of gaze is a result of breaking open the normal visual feedback loop, in which a corrective eye movement immediately eliminates (or minimizes) visual error in the retinal input. Note also how the gain of the saccades (i.e. their size) increases slightly within a given sequence of staircase saccades. This is a consequence of the known high gain of the oculomotor system, especially when feedback loops are broken (58).

**S1 Figure.**
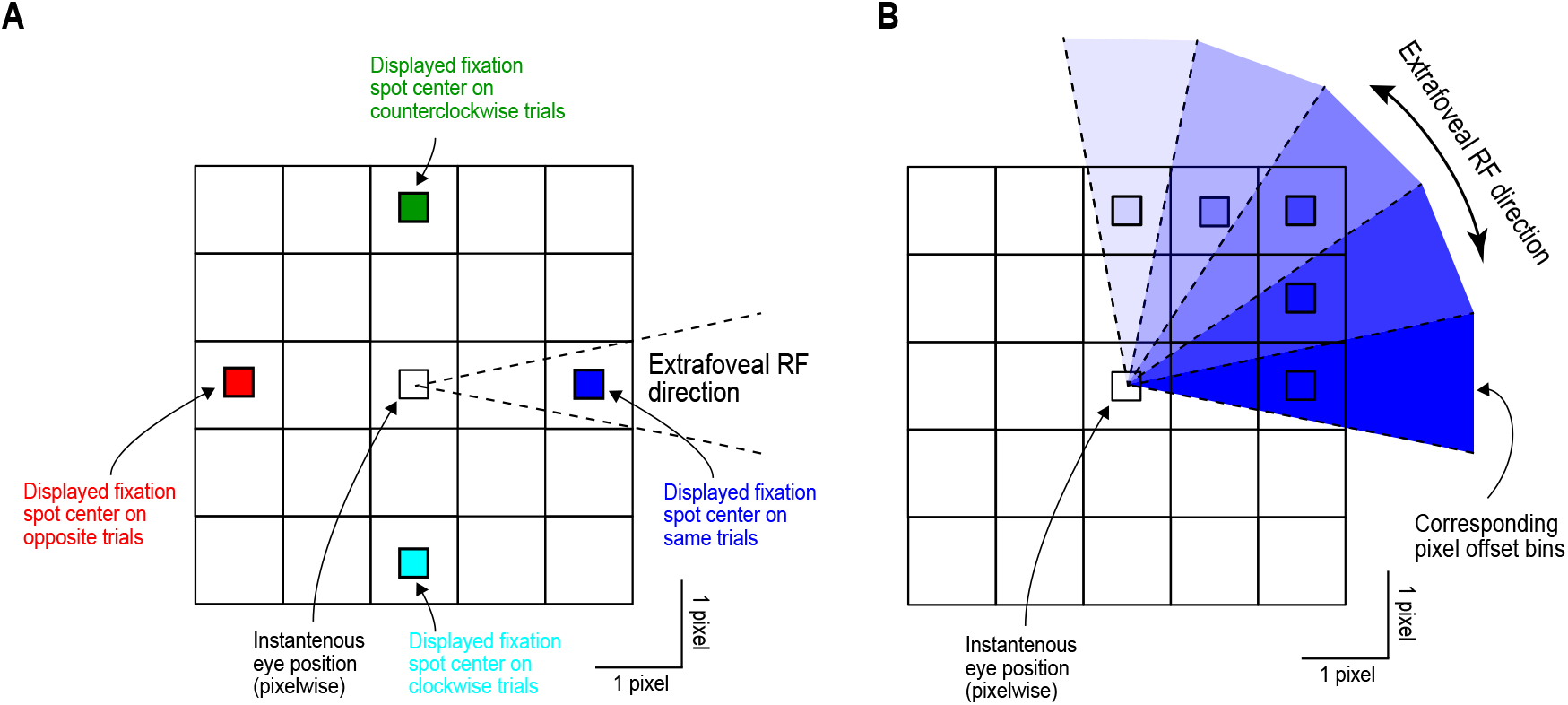
Implementing foveal retinal image stabilization under the constraints of a pixelated display. **(A)** During fixation, we measured instantaneous eye position. To force a constant foveal visual error, we centered the fixation spot at a position two pixels away from the instantaneous eye position (each square in the grid represents one single pixel). In the shown example, the extrafoveal RF was on the horizontal axis and to the right of gaze. Thus, a pixel offset to the right of instantaneous eye position was in the same direction as the upcoming extrafoveal visual stimulus (blue). In other trials, the fixation spot was centered two pixels away from instantaneous eye position, but this time in a direction directly opposite from the extrafoveal RF direction (red). In yet other trials, the forced foveal visual error was orthogonal to the direction of the extrafoveal RF (cyan and green). **(B)** Because extrafoveal RF direction could be completely arbitrary across recording sessions, we binned all possible RF directions into angular bins of 22.5 deg each. Thus, in the example of **A**, if the RF direction was within +/-11.25 deg in direction from the horizontal axis, we considered it close enough to being purely horizontal. For non-cardinal extrafoveal RF directions, we divided the full 360 deg range into 16 equal direction bins. Depending on which bin the extrafoveal RF direction fell in, the designated pixel offset on “same” trials was chosen accordingly, always from the square of possible positions 2 pixels away from instantaneous eye position but according to the directional bin (shaded regions indicate how each pixel offset position related to a corresponding extrafoveal RF direction bin). For simplicity, this panel shows examples from only one quadrant, and for only the “same” condition. The logic for all other quadrants applied similarly. Also, for “opposite” and “orthogonal” conditions, the bin 180 deg (for opposite) and +/-90 deg (for orthogonal) from the RF direction dictated the pixel offset position. Note that this approach meant slightly different foveal visual error amplitudes for different RF directions (e.g. compare 45 deg RF directions and 0 or 90 deg RF directions). This is unavoidable with pixelated displays, but the difference in foveal visual error amplitude was always extremely small in relation to the extrafoveal eccentricities that we tested (see S4 Figure). Also note that we could have used a smaller pixel offset of only 1 instead of 2 to alleviate this difference with angular bins; however, this would have severely hampered our angular resolution for representing arbitrary RF directions.

**S2 Figure.**
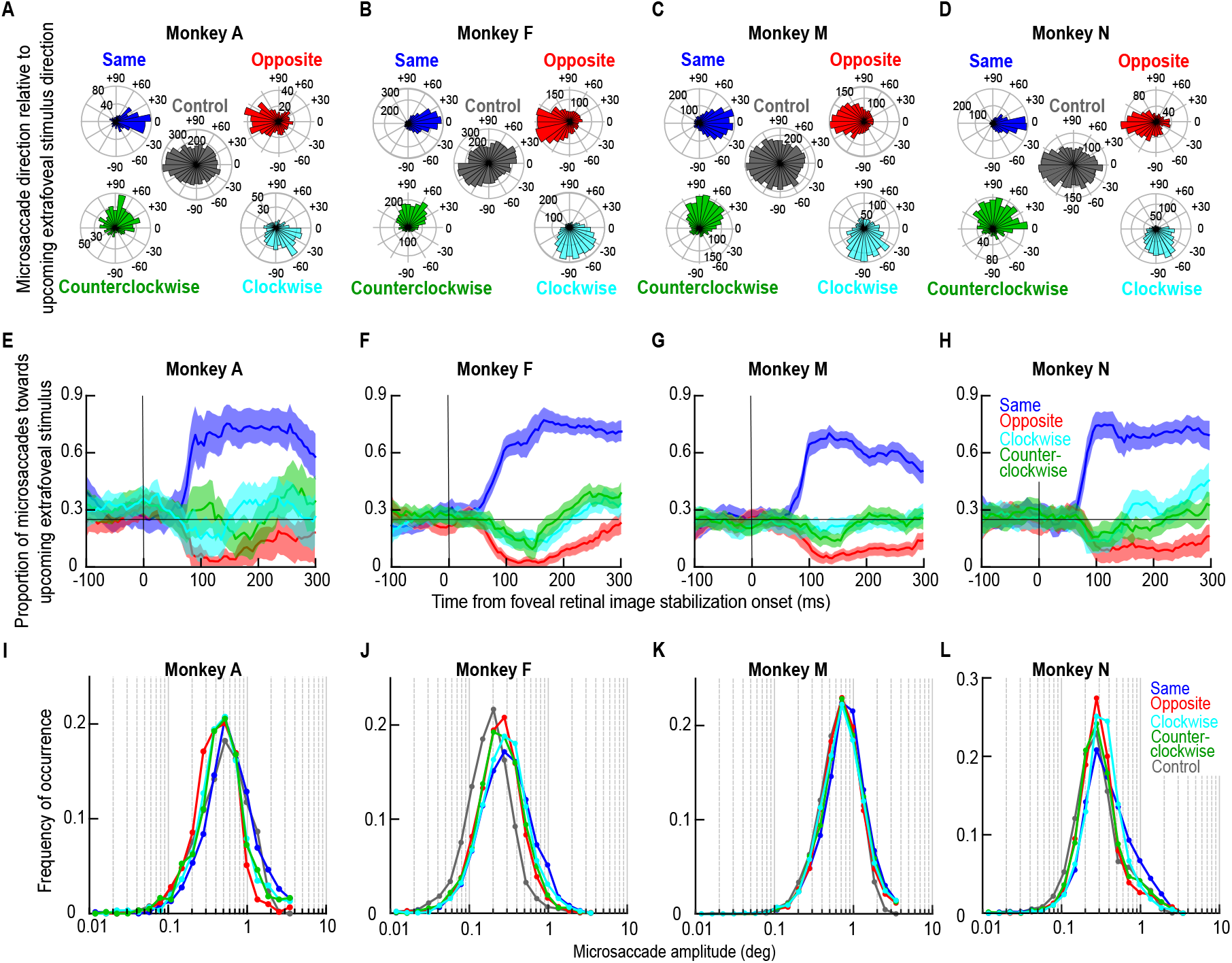
Individual monkey results from Fig. 1. **(A-D)** Same as Fig. 1C but for each monkey individually. In each case, we successfully controlled microsaccade directions using foveal retinal image stabilization. Note how for each monkey, the variance in microsaccade directions was lower in the same condition than in other conditions. This observation is further analyzed in S3 Figure. **(E-H)** Same as Fig. 1D but for each monkey individually. Shortly after the onset of retinal image stabilization, each monkey’s microsaccades were strongly biased by the direction of foveal visual error. The link between direction time courses and microsaccade rate evolution are also further explored in S3 Figure. **(I-L)** Same as Fig. 1F but for each monkey individually. In each case, the amplitude distributions of microsaccades were generally similar across all experimental conditions. Microsaccades were slightly larger, on average, under retinal image stabilization conditions than in control. This is expected since the average eye position error with successful fixation in control was expected to be near zero; however, with retinal image stabilization, we always forced a non-zero foveal error. Mean +/- SD microsaccade amplitudes in each monkey and condition were as follows. Monkey A: 35.5+/-23.7 min arc for control, 45.2+/-38.91 min arc for same, 28.9+/-21.9 min arc for opposite, 36.2+/-32.5 min arc for clockwise, and 36.9+/-33.9 min arc for counterclockwise; monkey F: 13+/-10.7 min arc for control, 22.8+/-21.1 min arc for same, 17.7+/-15.2 min arc for opposite, 21+/-19.5 min arc for clockwise, and 18.7+/-18.5 min arc for counterclockwise; monkey M: 44.2+/-23.5 min arc for control, 54.8+/-35.2 min arc for same, 49.9+/-32.1 min arc for opposite, 51.3+/-34.9 min arc for clockwise, and 52.2+/-34.4 min arc for counterclockwise; monkey N: 21+/-17.2 min arc for control, 28.8+/-23.5 min arc for same, 22.7+/-21.8 min arc for opposite, 24.6+/-23.6 min arc for clockwise, and 23.2+/-20.7 min arc for counterclockwise.

**S3 Figure.**
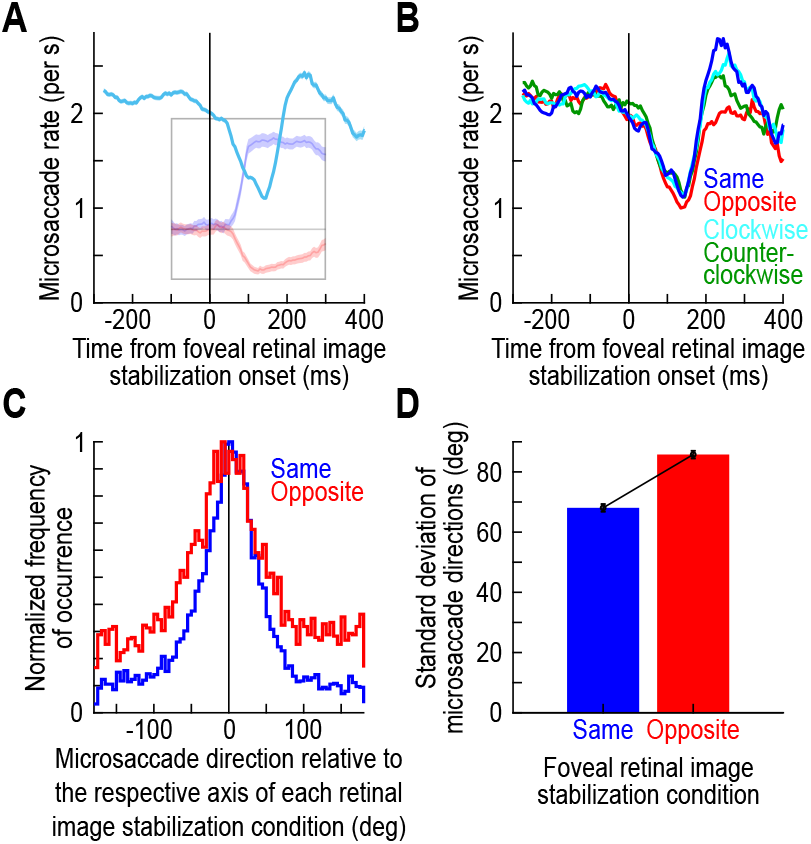
Further modulations of microsaccades caused by foveal retinal image stabilization. **(A)** Microsaccade rate (across all monkeys) around the time of onset of foveal retinal image stabilization. There was microsaccadic inhibition followed by rebound. The inhibition strength was weaker than previously observed with monkeys (e.g. refs. (14, 112)) likely due to the subtle retinal error that we introduced. Error bars denote SEM across trials. The faint inset shows the results of microsaccade direction time courses (Fig. 1D) to demonstrate the temporal relationship between microsaccade rate and direction modulations. An expected congruence between microsaccadic inhibition and direction biases was observed (8, 44). **(B)** When we split the rate curves according to the retinal image stabilization condition, we found that all retinal image stabilization conditions gave rise to the same time of microsaccadic inhibition. This means that any arousal signal caused by retinal image stabilization was independent of the direction of the foveal error. In the rebound phase, however, there were more microsaccades in the same condition than in the opposite and orthogonal conditions. This suggests a top-down influence (9) due to knowledge of the target location. **(C)** There was other evidence of top-down influences on the microsaccades. Specifically, we plotted the distribution of microsaccade directions relative to the axis of the retinal image stabilization condition. That is, for the same condition, 0 deg on the x-axis was the direction of the upcoming extrafoveal visual stimulus; for the opposite condition, 0 deg on the x-axis was the direction opposite the upcoming extrafoveal visual stimulus. Then, we plotted the distribution of microsaccade directions relative to the respective axis of retinal image stabilization. The y-axis range was normalized to the peak of each histogram, in order to highlight the variance difference between the two distributions. Microsaccade directions were less variable in the same condition than in the opposite condition. **(D)** Each bar shows the standard deviation of the respective distribution in **C**. The error bars denote 95% confidence intervals, based on bootstrapping (Methods). Microsaccade directions were less variable in the same condition. The underlying data are included in S1 Data.

**S4 Figure.**
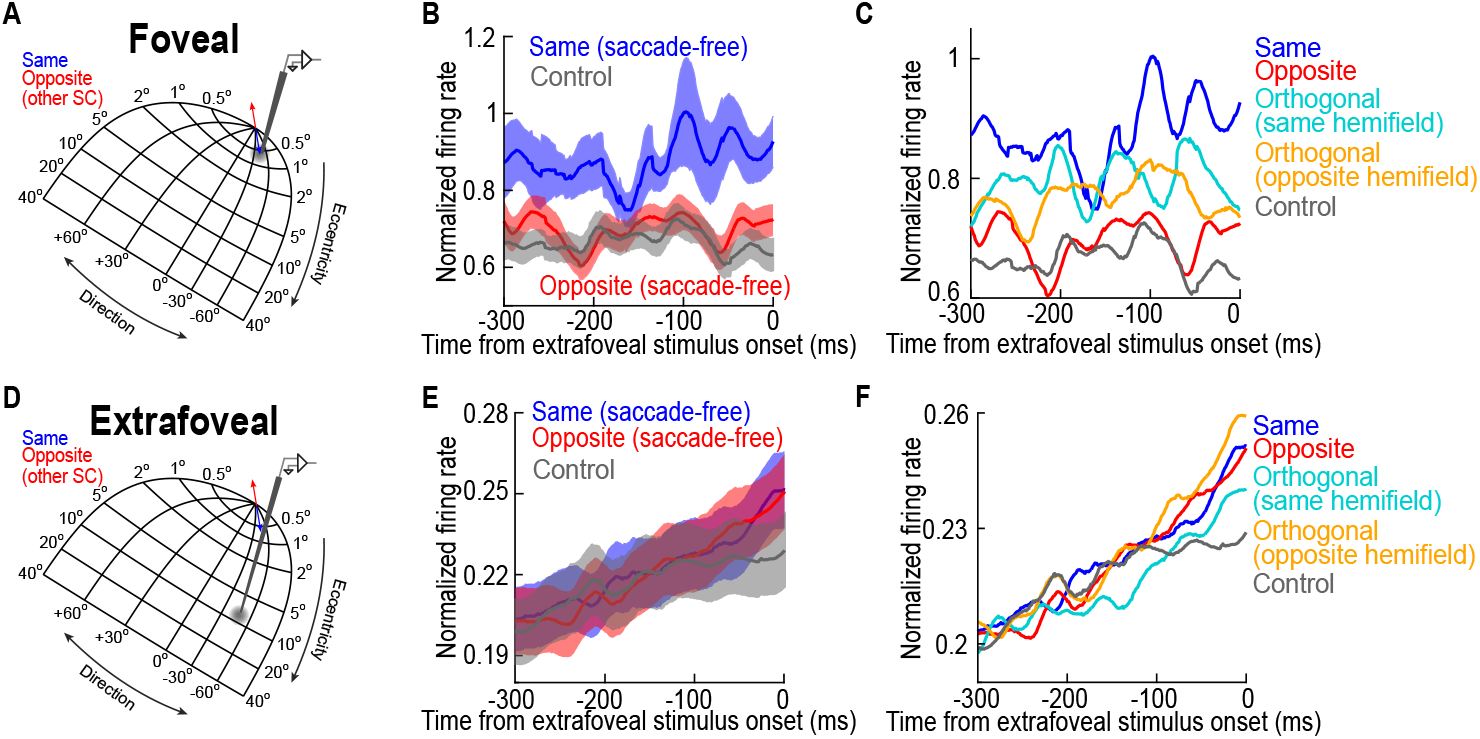
Exclusive differential modulation of foveal, but not extrafoveal, neuronal SC state by foveal retinal image stabilization; demonstration of the results obtained with all experimental conditions. **(A)** In a subset of experiments, we recorded from foveal SC neurons during our main behavioral task of our monkeys. **(B)** This panel is the same as that in Fig. 2B, except that we now also include the control condition. As can be seen, foveal retinal image stabilization had a clear directional modulation in the foveal SC neurons, explicitly elevating saccade-free activity in neurons congruent with the direction of the imposed foveal visual error direction (**A**). On trials with imposed visual error activating the other SC (opposite) or without experimental control over foveal visual error direction (control), average saccade-free activity was lower. Error bars denote SEM. **(C)** Same as **B** but now showing the orthogonal retinal image stabilization conditions as well. Note that here, unlike in the behavioral analyses (e.g. Fig. 1), we labeled the clockwise and counterclockwise retinal image stabilization conditions according to whether the foveal error direction was still in the same hemifield as the recorded neurons’ directions or whether it was in the opposite hemifield. Also note that we did not plot error bars in this panel to avoid visual clutter. As expected, average saccade-free activity on orthogonal stabilization trials was intermediate between that in the same and opposite trials. This expected because population coding in the SC (60), including in the foveal zone (52), suggests that an orthogonal foveal error direction might still partially activate the recorded foveal neurons. **(D)** We also checked pre-stimulus activity in the extrafoveal neurons of our database. **(E)** Same as Fig. 2F but now including the control condition. As expected, pre-stimulus extrafoveal SC activity gradually increased in anticipation of extrafoveal stimulus onset. This increase was slightly higher on retinal image stabilization trials than on control trials. This is because the monkeys could learn the statistics of the duration of retinal image stabilization (also see S3A Figure); thus, their expectation of extrafoveal stimulus onset was elevated. Nonetheless, there was no differential modulation on same and opposite trials, suggesting that the elevation was limited to target expectation and not due to differential modulation of foveal SC state. **(F)** In fact, there was no differential modulation of extrafoveal pre-stimulus activity by any of our four directions of retinal image stabilization (4 colors other than control). Once again, we did not plot error bars in this panel to avoid visual clutter. However, we statistically confirmed a lack of modulation of pre-stimulus extrafoveal neuronal activity by condition (one-way ANOVA across all five shown conditions; df=4; F=0.12; p=0.9765).

**S5 Figure.**
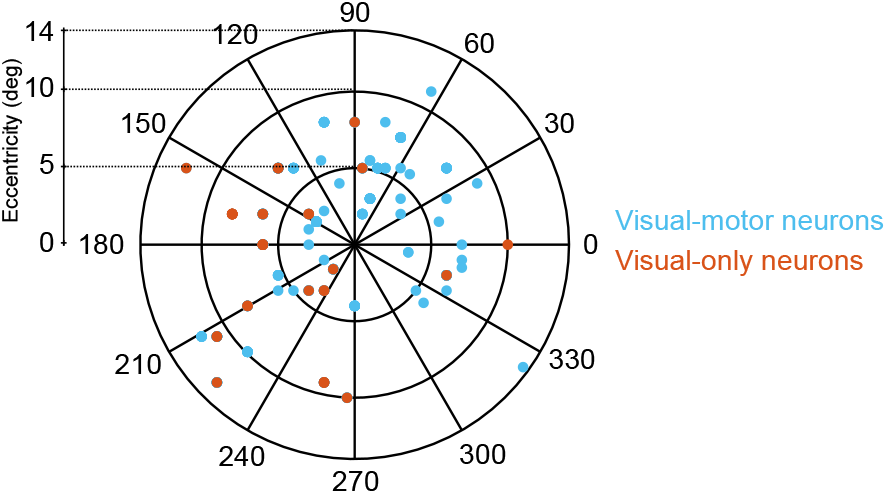
Visual response field hotspot locations for all extrafoveal neurons in our database. Our experiments spanned all four quadrants of visual space. We included 361 visual-motor neurons and 39 visual-only neurons. In all cases, there was no microsaccade-related activity in any of the neurons. Thus, the neurons’ response fields did not extend foveally into the SC region that was modulated by foveal retinal image stabilization during the pre-stimulus interval. Consistent with this, there was no differential modulation of these neurons’ pre-stimulus activity by the direction of foveal retinal image stabilization (Fig. 2, S4 Figure). Also note that multiple neurons recorded within a single session had overlapping response field locations. Thus, we used a single stimulus location for such neurons per session. As a result, this figure, plotting the stimulus location of each session as an individual symbol, contains fewer than 400 symbols.

**S6 Figure.**
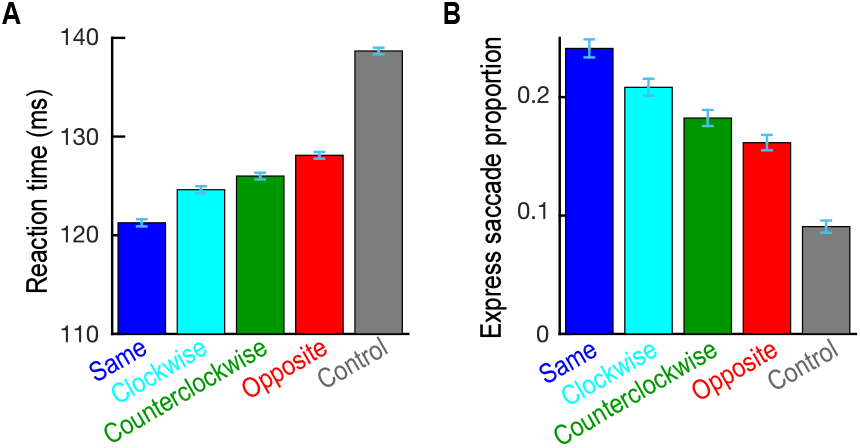
Orthogonal retinal image stabilization directions always resulted in intermediate results between the same and opposite conditions, consistent with the microsaccadic directional modulations of Fig. 1, as well as the foveal neuronal modulations of Fig. 2A-C, S4A-C Figure. **(A)** Similar to Fig. 2H (left), but now additionally including the clockwise and counterclockwise foveal retinal image stabilization conditions, as well as the control condition. All foveal retinal image stabilization conditions were associated with shorter saccadic reaction times than in the control condition. This is due to the expectation of extrafoveal stimulus onset, and the fact that retinal image stabilization onset increased the likelihood of upcoming extrafoveal stimulus onset (Methods and S4D-F Figure; also see S3B-D Figure). However, even though pre-stimulus activity was the same across all four directions of foveal retinal image stabilization (Fig. 2E-G, S4D-F Figure), reaction time clearly depended on the direction of forced foveal visual error (df=4, F=355.63, p=0; one-way ANOVA across all five conditions): same trials had the fastest reaction times, opposite trials had the shortest reaction times, and orthogonal trials gave intermediate results. Error bars denote SEM. Identical conclusions were also reached from each monkey individually (see S8 Figure). **(B)** Consistent with **A**, the likelihood of express saccades was higher in retinal image stabilization than in control trials. However, once again, even though pre-stimulus extrafoveal SC activity was similar across all four directions of forced foveal visual error (Fig. 2E-G, S4D-F Figure), express saccade likelihood followed the differential modulation of foveal SC activity (S4A-C Figure). These results indicate that the influence of foveal state was mediated through modulating visually-evoked extrafoveal activity (Figs. 3–5). Error bars denote 95% confidence intervals. The underlying data are included in S1 Data.

**S7 Figure.**
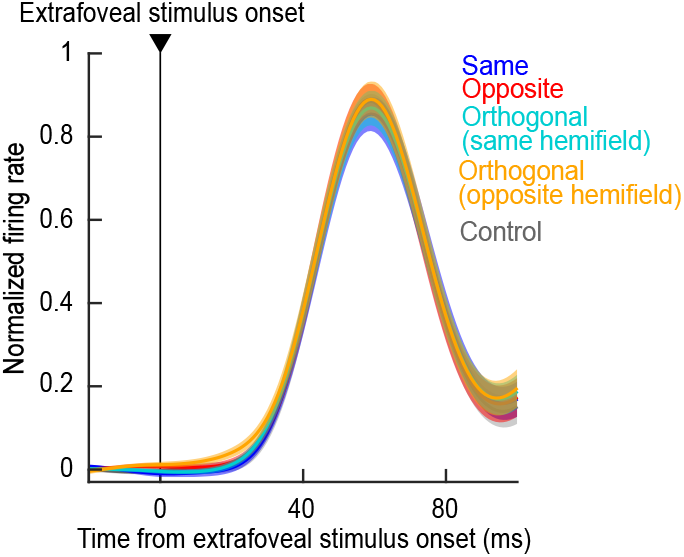
Lack of modulation in the sensitivity of extrafoveal visual-only SC neurons by experimental control over foveal oculomotor state. Averaged, normalized, and baseline-subtracted firing rates in the control, same, opposite, and two orthogonal foveal retinal image stabilization conditions. Unlike visual-motor neurons (Fig. 4), our population of visual-only neurons (39 neurons) showed no differences in extrafoveal visual sensitivity as a function of forced foveal visual error directions. Error bars denote SEM.

**S8 Figure.**
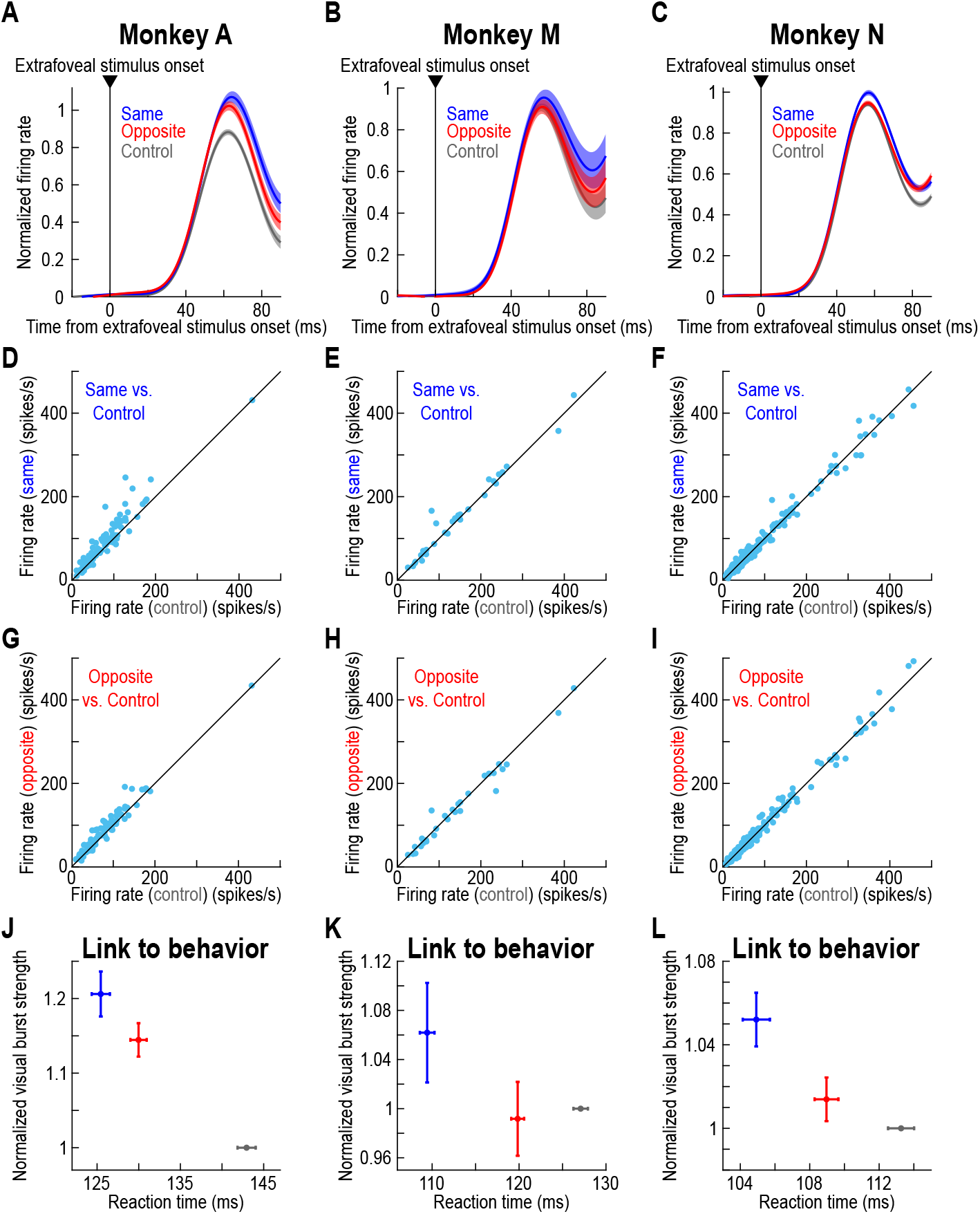
Individual monkey neuronal and behavioral results. **(A-C)** Like Fig. 4A, but for each monkey individually. Error bars denote SEM. **(D-F)** Like Fig. 4B, but for each monkey individually. **(G-I)** Like Fig. 4C, but for each monkey individually. **(J-L)** Like Fig. 4E, but for each monkey individually. All monkeys showed consistent results, which were also consistent with other studies (31, 34). The underlying data are included in S1 Data. All other conventions like in Fig. 4.

**S9 Figure.**
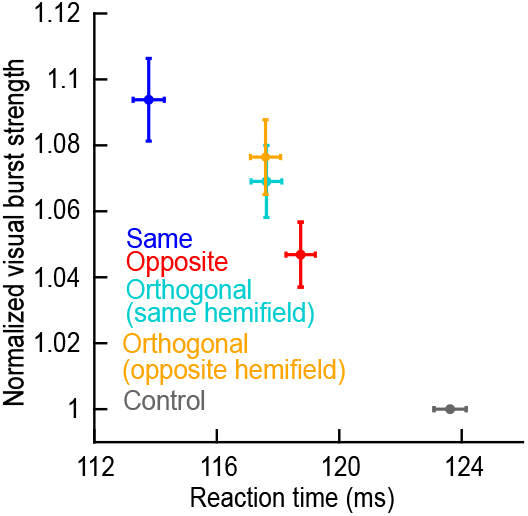
Intermediate results for orthogonal foveal retinal image stabilization conditions in both behavior and extrafoveal SC visual sensitivity. For our visual-motor neurons (361 neurons), we plotted visual burst strength as a function of reaction time in control, as well as in the four foveal retinal image stabilization conditions. Consistent with Fig. 4, S6 Figure, foveal retinal image stabilization was associated with faster reaction times than in control. This is due to target expectation (e.g. S4E-G Figure). Nonetheless, reaction time was differentially modulated by foveal oculomotor state (also see S6 Figure), and this was directly evident in extrafoveal SC visual sensitivity (similar to Fig. 4 but now including the orthogonal foveal retinal image stabilization conditions as well). Thus, exclusive differential modulation of foveal, but not extrafoveal, oculomotor and pre-stimulus SC state (Figs. 1, 2, S4 Figure) was sufficient to also cause differential modulation of extrafoveal visual sensitivity (y-axis) and, ultimately, behavior (x-axis). All other conventions are like in Fig. 4. The underlying data are included in S1 Data.

**S10 Figure.**
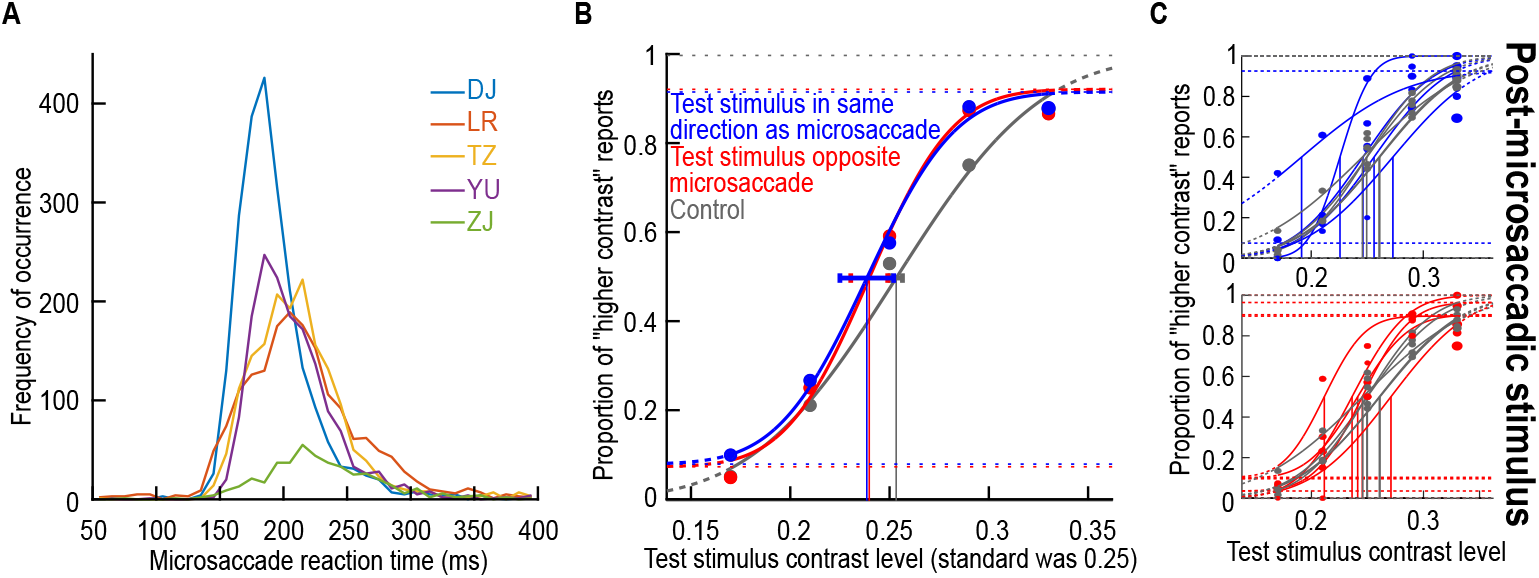
Similar influences of voluntary human microsaccade generation on extrafoveal contrast sensitivity. **(A)** Distribution of microsaccade reaction times relative to fixation spot jump in our main human experiment of Fig. 6 (Methods). All five human subjects (different colors) successfully generated microsaccades with expected reaction times after fixation spot jumps. **(B, C)** Similar to Fig. 6B, C but now for test stimuli appearing within 50 ms after, rather than before, microsaccade onset. Relative to control, there was evidence of slightly enhanced sensitivity in both the same (p=0.005, permutation test; Methods) and opposite (p=0.0151, permutation test; Methods) conditions. However, unlike for pre-microsaccadic stimulus onsets (Fig. 6B, C), there was clearly no difference between the same and opposite conditions. Thus, the differential, direction-dependent modulation by microsaccades was restricted to the pre-movement epochs (Fig. 6) (30). Error bars denote SEM across 5 subjects.

**S11 Figure.**
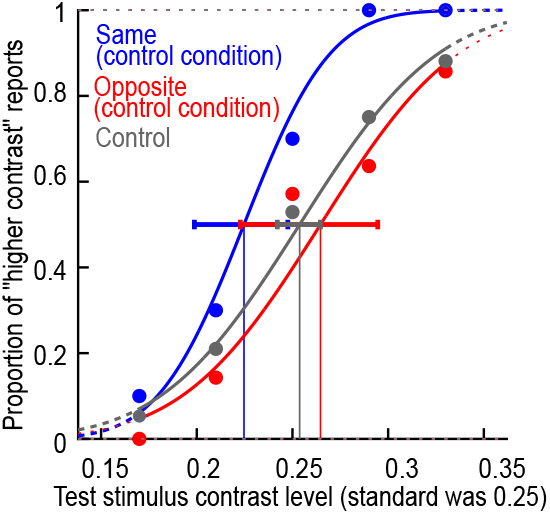
Similar pre-microsaccadic modulations even in control trials. By classifying microsaccade directions in the control condition, we observed similar results to Fig. 6B, C. When test stimuli appeared within 50 ms before microsaccade onset and the microsaccade was in the direction of these stimuli (same), there was enhanced contrast sensitivity relative to trials without microsaccades (control). For opposite microsaccades, there was slightly reduced contrast sensitivity. Naturally, due to lack of experimental control over the directions of peri-stimulus microsaccades in this condition, we had fewer trials with proper directions and times than in Fig. 6B, C. However, just like in Figs. 2D, 5, when the right combination of microsaccade directions and times occurred, the same trends were clearly in play also during control trials. This is also consistent with our prior work (4, 30, 32). Error bars denote 95% confidence intervals.

**S12 Figure.**
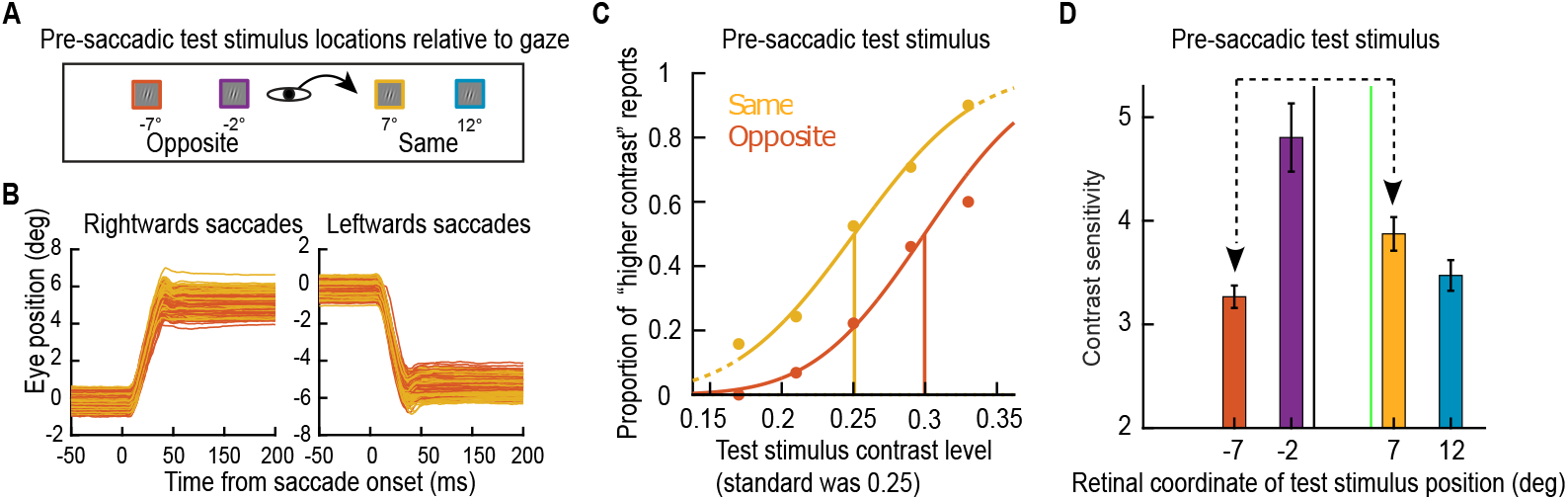
Pre-movement enhancement of contrast sensitivity also ahead of large-saccade endpoints, not just microsaccades. **(A)** We asked human participants to make 5 deg saccades to the left or right. The experiment was otherwise identical to that of Fig. 6. That is, before the saccade instruction, there were standard stimuli. Then, there was a saccade instruction (this time requesting 5 deg saccades). At a random time, a test stimulus was presented at 2 deg, 7 deg, or 12 deg from the fixation spot location, and we analyzed trials in which this stimulus appeared within 50 ms before saccade onset. This way, the stimuli at 7 deg on either side of fixation were at retinotopically matched positions, with the only difference being that one of them was ahead of the saccade target endpoint and in the same hemifield (same direction), whereas the other was in the opposite hemifield. **(B)** Sample subject result showing no difference between left and rightwards saccades when we presented pre-saccadic test stimuli at 7 deg (either in the same direction as the 5 deg saccade, orange, or in the opposite direction, red). **(C)** Sample subject contrast sensitivity curves showing increased sensitivity ahead of the saccade target when the test stimulus was in the same direction relative to when it was in the opposite direction. In both cases, the test stimulus appeared within 50 ms before saccade onset, and it was at the same retinotopic eccentricity (either in the same or opposite hemifield from the saccade target location). Vertical lines indicate the threshold contrasts in each condition. **(D)** Average contrast sensitivity measures (calculated as the inverse of perceptual thresholds obtained from psychometric curves like those in **C**) across eight human subjects. The data are for pre-saccadic target onsets (Methods). The green vertical line marks the 5 deg saccade eccentricity. The dashed line indicates that the same and opposite conditions shared the same retinotopic coordinates on the retina (since the stimuli appeared before saccade onset). Despite that, contrast sensitivity was higher in the same direction, and this was true even though the test stimulus was more eccentric than the saccade amplitude itself (similar in principle to our microsaccadic results). Even at 12 deg, contrast sensitivity in the same direction was not lower than contrast sensitivity at 7 deg in the opposite direction, despite being 7 deg ahead of the saccade endpoint (instead of only 2 deg ahead of it). Statistically, pre-saccadic performance at 2 deg was the best because of proximity to the fovea (p<0.0001; permutation test; Methods). Performance at 7 and 12 deg in the same direction as the saccade was higher than performance at 7 deg in the opposite direction, even though both 7 and 12 deg were ahead of the saccade target at 5 deg (7 deg: p<0.0001; permutation test; 12 deg: p=0.0484; permutation test). Error bars denote SEM across 8 subjects. The underlying data are included in S1 Data.

## Notes

### Competing Interest Statement

The authors have declared no competing interest.

### Summary of Updates

- Updated analysis of Fig. 2 to include genuine opposite and towards movements - Updated Fig. 5 as above - Did further analyses of retinal image stabilization effects - Added individual monkey results - Performed final journal formatting requirements

